# Weak but consistent genomic signals of intragenerational selection during estuarine migration in the European eel (*Anguilla anguilla*)

**DOI:** 10.64898/2026.02.01.703082

**Authors:** Stellia Sebihi, Aurélie Manicki, Christophe Klopp, Mathieu Gautier, Pascale Coste, Emmanuel Huchet, Maren Ortiz-Zarragoitia, Valérie Bolliet, Sylvie Oddou-Muratorio

**Affiliations:** Université de Pau et des Pays de l’Adour. E2S UPPA. INRAE. ECOBIOP. MIRA. UMR 1224 – St-Pée-sur-Nivelle. France; CBET Research Group. University of the Basque Country – Plentzia, Basque Country, Spain; INRAE, UMR Ecobiop (INRAE—UPPA) – St-Pée-sur-Nivelle. France, France; Sigenae, Genotoul Bioinfo, BioInfoMics, MIAT UR875, INRAE – Castanet Tolosan, France; INRAE, UMR CBGP (INRAE—IRD—Cirad – Montpellier SupAgro) – Montferrier-sur-Lez, France

**Keywords:** Glass eel, Selection, Spatial sorting, Genetics, Pool sequencing, Panmixia, Migration, *Anguilla anguilla*

## Abstract

Migration events can act as strong selective filters by spatially sorting individuals according to their migration ability, behaviour, and associated functional traits. The European eel, a panmictic and threatened fish, presents various estuarine migration patterns at juvenile stage (glass eel), ranging from sedentarization in brackish/saltwater of the estuary (non-migrant phenotype) to upstream colonisation of freshwater ecosystems (migrant phenotype). We hypothesize that migration propensity is partly genetically determined in glass eel, and that migration-related genotypes are spatially sorted during estuarine migration. To test these hypotheses, we first collected six pools of individuals over three years at two extreme sites along a gradient from ocean to Adour River tidal limit (Ocean vs. Upstream). Secondly, we collected additional glass eels and phenotypically sorted migrant vs. non-migrant individuals using an experimental device mimicking alternating tidal currents, producing two other pools. Whole genome pool sequencing and analysis of these eight pools generated 18.99 10^6^ SNP variants.

Controlling for linked selection through a local score approach, we found five best outlier SNPs with a significant genetic differentiation between Ocean *vs.* Upstream sites (average *F_ST_* = 0.21) compared to the pangenomic estimate (*F_ST_* = 0.0086). These five SNPs were all found in the same gene (gpb2), involved in interferon-mediated antiviral immune responses. We also found 28 best outlier SNPs with a significant genetic differentiation between migrant *vs.* non-migrant phenotypes (average *F_ST_* = 0.51). They were located in genes mainly involved in neuronal development, cell migration and tissue remodelling, transcriptional regulation, and metabolic or stress-related processes.

Our results support that variation in eel migration propensity is partly genetically determined and that, while panmixia maintains high level of genetic diversity, spatial sorting could promote intra-generational genetic divergence between habitats of European eels. However, the absence of shared genes among the best outliers between *in-situ* and experimental contrasts suggests a complex and context-dependent genetic control of migration.

## Introduction

Migration, defined as the movement of organisms among two (or more) locations is a critical component of many animal life cycles, during which individuals face abrupt environmental changes and strong physiological constraints (Dingle, 2014). Migration events can therefore act as strong selective filters by spatially sorting individuals according to their migration ability, behaviour, and associated functional traits. When variation in these traits has a genetic basis, such sorting is expected to generate intragenerational phenotypic and allelic shifts, reflecting a spatial selection differential rather than an evolutionary response *sensu stricto*. In species with panmictic reproduction, where alleles are reshuffled at each generation across the distribution range, this spatial selection differential is not expected to result in classical local adaptation, as alleles that are advantageous in one environment may be maladaptive in another (Bernatchez, 2016; Babin et al., 2017). However, theory and empirical studies have shown that spatially varying selection can maintain adaptive polymorphism in panmictic populations by sorting locally favoured alleles within each generation (Gagnaire et al., 2012). Although alleles under selection are often transient and may ultimately reach fixation if selective pressures are asymmetric, this evolutionary mechanism referred to as protected polymorphism model (Levene, 1953), is a pertinent conceptual framework to study adaptation in a panmictic species.

The European eel (*Anguilla anguilla*) provides an exemplary system to investigate this issue. Allegedly breeding in the ocean as a single panmictic population (Schmidt, 1923), the European eel spends most of its life cycle in a wide range of brackish to freshwater habitats. A critical stage occurs after the long-distance transoceanic migration of larvae to the continental slope, when juveniles, called glass eels, potentially colonise freshwaters to grow (Tabeta & Mochioka, 2003; Briand et al., 2005). At this stage, some individuals stop their migration and settle in brackish waters within the estuaries, others migrate further and settle in freshwater habitats, while other ones move back and forth between these habitats (Tsukamoto, Nakai & Tesch, 1998; Daverat et al., 2006). These estuarine migration patterns have major consequences for the population dynamics of European eel, which is classified as critically endangered by the IUCN. Indeed, glass eels that colonise watershed develop mainly as females, whereas those that settle in estuaries tend to produce males (Davey & Blaxter, 2010; Geffroy & Bardonnet, 2016), with different rates of individual growth and density in each habitat. It is therefore crucial to investigate the determinism of glass eel migration to better understand how the spatial sorting of individuals during this critical stage affects the sex ratio and demographic dynamics of this emblematic and threatened species.

Interindividual variation in glass eels’ propensity to migrate within the estuaries may arise from differences in physiological capacities and/or decision-making processes that affect migratory behaviour, these variations being possibly shaped by genetic and/or environmental factors, and their interactions. Several studies have investigated the hypothesis that estuarine migration propensity is conditional, i.e. based on individual energy status (Edeline, Dufour & Elie, 2005; Bureau Du Colombier et al., 2007; Edeline, 2007; McCleave & Edeline, 2009; Liu et al., 2019). Most glass eels do not feed during migration (Bardonnet & Riera, 2005), relying on energy reserves accumulated during oceanic migration to sustain activity and reach freshwater (Tesch & Thorpe, 2003). In line with the conditional hypothesis, some studies showed that glass eels with high energy reserves had a greater propensity to migrate than those with low reserves (Edeline et al., 2004; Bureau Du Colombier et al., 2007; McCleave & Edeline, 2009). However, other experimental studies in European or American eels (*Anguilla rostrata*) found that lipid contents or body condition had no effect on glass eel’s propensity to migrate or on salinity preferences (Edeline, Dufour & Elie, 2005; Bureau Du Colombier et al., 2007; Boivin et al., 2015; Gaillard et al., 2015; Bolliet et al., 2017; Liu et al., 2022).

By contrast, whether estuarine migration propensity has a genetic basis remains largely unknown, partly because research on eels has long focused on the consequences of panmixia (Pujolar, Jacobsen & Bertolini, 2022), or more recently on oceanic migration genetic determinism (Liu et al., 2024). The eels’ unique reproductive strategy, which includes panmixia and a large effective population size, is known to result in high genetic diversity and low overall genetic differentiation between growth habitats at large spatial scales (Gagnaire et al., 2012; Pujolar et al., 2014; Ulrik et al., 2014). Nonetheless, several studies have found evidence of spatially varying intragenerational selection among growth habitats in *Anguilla* species. For example, Gagnaire et al. (2012) found 13 Single Nucleotide Polymorphism (SNP) loci significantly correlated with temperature across a latitudinal gradient in the American eel. Their study introduced the idea that habitat-specific adaptive alleles could be maintained over generations by spatially variable selection despite panmixia. Similarly, Pujolar et al. (2014) reported high genetic differentiation driven in particular by temperature differences along a latitudinal gradient in European eels. However, these studies conducted over regional to larger spatial scales did not account for the possible local variation among salt- and freshwater habitats at estuarine scale. More closely related to estuarine migration propensity, Pavey et al. (2015) found 331 co-varying loci associated with the divergence between freshwater and saltwater ecotypes in American eels. They proposed two mechanisms to explain this divergence, *i.e.*, genotype-dependent habitat choice and intragenerational spatially variable selection. However, and because they studied individuals at a late stage of life-history, they were unable to disentangle whether these mechanisms operate throughout an individual’s life or at a specific life stage.

Our hypotheses are that the propensity to migrate within the estuary is partially genetically determined in European eel, and subject to spatially varying selection. To address previous limitations, we focus on the critical stage of glass eel estuarine migration. First, we analyse *in-situ* genomic differentiation between two sites: “Ocean” located at the mouth of the Adour estuary (saltwater) *vs.* “Upstream” located 40 km from the estuary mouth (freshwater). We hypothesize that the Ocean site hosts a mixture of migrant glass eels (individuals that will continue their migration to upstream site) and non-migrant glass eels (individuals that settle there), whereas the Upstream site hosts mainly migrant glass eels. Hence, we predict allele frequency differences between sites at the genes related to propensity to migrate in the estuary or habitat choice (due to spatial sorting). These sites were sampled in three different years, capturing three independent glass eels’ cohorts. Secondly, based on additional sampling of glass eels, we use an experimental approach where individuals are phenotypically sorted according to their propensity to swim and synchronize their swimming activity with the change in water current direction (mimicking an alternating tide). We hypothesize that this behaviour is a main component of migration propensity: indeed, glass eels migrate up estuaries using selective flood transport i.e., they swim in the water column during the flood and move towards the substratum during ebb tide (Jellyman, 1979; Forward & Tankersley, 2001). Hence, we predict allele frequency differences between extreme phenotypes at their causal genes (*i.e.*, phenotypic sorting). We use a whole-genome pool-sequencing approach (Schlötterer et al., 2014; Nunez et al., 2025), pooling DNA from approximately 35 individuals at each site for the *in-situ* approach (Ocean *vs.* Upstream, repeated over three years, amounting to six pools) or in each phenotypic group in the experimental approach (two pools). We then employ state-of-the-art population genomic approaches to identify loci with significant differentiation (*i.e., F_ST_* outliers) between sites or phenotypic groups (Gautier, 2015; Olazcuaga et al., 2021). Combining these *in-situ* and experimental approaches allows us to pinpoint the genomic regions potentially involved in the propensity to migrate, and provides valuable insights into the biological mechanisms driving estuarine migration of European glass eels.

## Material and methods

### Ethics

The procedures performed during experimentation in a controlled environment were validated by the ethics committee N°073 “Aquitaine Poissons-Oiseaux” (reference: APAFIS-202207 1416031365). All this experiment was performed in compliance with the EU legal frameworks, specifically those relating to the protection of animals used for scientific purposes (*i.e.,* Directive. 2010/63/EU), and under the French legislation governing the ethical treatment of animals (Decret no. 2013–118. February 1st. 2013).

### Study site and sampling design

#### In-situ approach: sampling downstream and upstream of the Adour estuary

Glass eels were sampled at two sites along the Adour estuary: the first site “Ocean” is located downstream from the mouth of the estuary at Moliets-et-Maa, in saltwater, and it is subject to dynamic tides. The second site “Upstream” is located at Josse, 40 km Upstream from the mouth of the Adour estuary, in freshwater, close to the tidal limit. Glass eels were sampled using a dip-net at night during black moon period at the end of November or beginning of December and during flood tide, targeting 80 individuals per site. Each site was sampled in 2019, 2020 and 2022, yielding three temporal cohorts considered as replicates, as glass eels arriving each year originate from independent panmictic spawning events.

After fishing, glass eels were transported to the laboratory and kept in a tank containing aerated water from the capture site. They were maintained at 12.0°C ± 0.5°C and in the dark for the rest of the night. In the next morning following the sampling, glass eels were anesthetized and killed using lethal dose of anesthesic. Individual initial length (±1.0mm) and wet weight (Sartorius CP 153 balance, ±1.0mg) were then measured and all glass eels were then stored at −20°C until analysis.

#### Experimental approach: phenotypic sorting in controlled conditions

At Moliets-et-Maa in November 2022, 180 additional glass eels were sampled as described above, brought to the laboratory and kept in a tank containing seawater from the capture site. They were maintained at 12°C ± 0.5°C with a photoperiod of 10 hours of light (0.2-0.3 *µ*W.cm²) and 14 hours of darkness in a continuously aerated tank (10L/14D). During the next 96 hours, the seawater was progressively diluted with freshwater (1/4 of the water the first two days, 1/3 on the other days).

After 96 h acclimatization, glass eels were tagged using Visible Implant Elastomer (combination of two or three hypodermic VIETag spots) and then released to wake up in an aerated tank. After a night of recovery, tagged individuals were transferred to two annular tanks (90 glass eels in each) exposed to a change in water current direction every 6.2 hours to mimic the tides as described in Liu et al. (2019). The water and room temperature were kept at 12.0 ± 0.5°C and fish were submitted to a 10L/14D photoperiod (0.2-0.3 *µ*W.cm² during photophase) and a constant UV light (0.6 *µ*W.cm²) to detect VIETag at night. A video recording of 15 seconds every 40 minutes over a period of five days was used to determine individual behaviour. The sampling session duration was chosen to allow fish to pass once through the camera’s field of view when swimming in the water column. A total of 107 sessions of 15 seconds were obtained for each batch.

To migrate up the estuary, glass eels synchronize their swimming activity with the tide using selective flood transport (circatidal rhythm; Jellyman, 1979; Forward & Tankersley, 2001). They swim with the current on flood tide and hide in the substratum on ebb tide. Therefore, to evaluate the propensity of glass eels to migrate under experimental conditions, we analysed the synchronization of their swimming activity to the change in water flow direction. Glass eels were considered to have a high propensity to migrate within the estuary when their swimming activity was synchronized with water current reversal. In contrast, glass eels that did not synchronize and mostly buried in the substrate regardless of the current direction were considered to have a low propensity to migrate. Out of the 180 individuals monitored, we selected the 40 displaying the most extreme behaviour in each category (*i.e.,* the most synchronized, called ExpMigrant and the least active, called ExpNonMigrant), which were then anesthetized, killed using lethal dose of anesthesic, and stored at −80°C after immersion in liquid nitrogen.

### Genomic data acquisition

#### DNA extraction

DNA was extracted from 1 cm of muscle tissue from the tail of each individual, that had been stored at −20°C since sampling. To avoid DNA degradation, muscle sampling was performed on ice. Fish muscle DNA was extracted with saturated NaCl and chloroform in 96-well plates. The extraction process involved lysing the tissue in 200 µL of extraction buffer (25mM EDTA pH 8, 1% SDS, 75mM NaCl) containing proteinase K 20 mg.ml^-1^ overnight at 55°C. Next, RNA was digested with 4µL of 100 mg.ml^-1^ RNase A during a 5-minute incubation at room temperature. Then, 100 *µ*L of a saturated 5M NaCl solution and 300 *µ*L of chloroform were added to each sample. After 10 minutes of incubation and inversion mixing at room temperature, the samples were centrifuged for 10 minutes at room temperature (2000 rpm). DNA precipitation was realized with 250 *µ*L of isopropanol and 2 *µ*L of linear acrylamide (5 mg.ml^-1^), then a mix step during 5 minutes at room temperature, then by a centrifugation at 3700 rpm for 5 minutes at 4°C. The DNA pellet was washed with 500 *µ*L of 70% ethanol, followed by a shaking step of 1 hour at 4°C and a centrifugation at 3700 rpm during 5 minutes at 4°C. Once the pellet had dried, it was resuspended in 50 *µ*L of 1X TE buffer. The quality and quantity of DNA extracted from each individual were assessed using a 1.5% agarose gel, coupled with analysis using the QUBIT 2.0 fluorometer (Thermo fisher Scientific) and the Nanodrop.

#### Pooled whole genome Sequencing (PoolSeq)

We grouped individuals into eight pools (Table S1), including (1) six pools for the *in-situ* approach (three pools Ocean2019, Ocean2020 and Ocean2022; three pools Uptream2019, Uptream2020 and Uptream2022) and (2) two pools for the experimental approach (ExpMigrant and ExpNonMigrant). To prepare the corresponding eight PoolSeq libraries, individual DNAs were pooled in equimolar ratios. DNA quantity and quality of each pool were assessed using a QUBIT 2.0 fluorometer to estimate concentration, and nanodrop analysis to determine 260/280 and 230/280 quality ratios (Table S1). Pooled DNA libraries were constructed using Illumina’s TruSeq Nano DNA HT library preparation kit, according to the manufacturer’s protocol. Sequencing was performed using Illumina NovaSeq X, targeting a sequencing depth of 50X and generating paired-end reads 150 bp in length. Library preparation and sequencing were performed at the GeT-Plage platform (INRAE, Toulouse, France), a member of the GeT Genotoul platform.

#### Bioinformatic analyses

Raw paired-end reads were filtered using fastp 0.23.2 with default parameters to remove adapter sequences and trim for low quality bases (*i.e.,* with a phred-quality score < 20). Filtered reads were then aligned to the NCBI reference genome of *A. anguilla* (fAngAng1.pri, NCBI) using default parameters of the mem command from bwa package version 0.7.17 (Li & Durbin, 2009). The resulting alignments were compressed, merged, and then indexed with the SAMtools package version 1.9 (Li et al., 2009) using default parameters.

Variant calling was performed using the Haplotype Caller implemented in FreeBayes (Garrison & Marth, 2012), which does not require realignment around indels and base recalibration, jointly considering all 8 Pool-Seq alignment files, with options “-K” (to output all alleles that pass input filters, regardless of genotyping outcome, assuming a pooled sequencing model); “-C 1 -F 0.01” (at least one count and a fraction of 1% of the counts of the alternate allele to evaluate the position); “-G 5” (at least 5 counts supporting an alternate allele in all sample to retain the allele); “-E −1” (to disable clumping of contiguous variants into complex alleles); “--limit-coverage 500” (to downsample per-sample coverage to 500 reads if greater than this coverage); “-n 4” (to evaluate only the best 4 alleles, ranked by sum of supporting quality scores), “-m 30 -q 20” (minimum mapping and base qualities set to 30 and 20, respectively). Finally, the resulting vcf file was annotated using the NCBI reference gene model for *A anguilla* (genome assembly GCF_013347855.1_fAngAng1.pri), provided as a GFF3 file, and SnpEff version 4.3T (Cingolani et al., 2012).

Read processing, alignment and variant calling were performed using the computational resources of the Genotoul bioinformatics platform (INRAE, Toulouse, France).

The final vcf file was transformed into a pooldata object using *vcf2pooldata* function of the poolfstat package (Gautier et al., 2022), with the options min.rc = 2, to exclude SNPs with a read count below 2 (indicative of too low coverage), and min.maf = 0.01, to filter out poorly polymorphic sites. An additional filtering step was then applied to remove SNPs with either excessively low or high coverage and insufficient polymorphism, using pooldata.subset with min.cov.per.pool = 20, cov.qthres.per.pool = c(0, 0.99), and min.maf = 0.05. This informative SNP dataset was used to compute genetic differentiation between pools, and to run the following outlier tests.

### *F_ST_* outliers tests of within-generation selection

To detect loci under selection from poolseq data, we employed two complementary methods: (i) *BAYPASS* (Gautier, 2015) which estimates the contrast statistic *C₂* to compare allele frequencies between groups defined by a binary covariate (e.g., Ocean vs. Upstream; Olazcuaga et al., 2021) and (ii) PCAdapt (Duforet-Frebourg et al., 2016; Luu, Bazin & Blum, 2017), a model-free approach based on Principal Component Analysis (PCA) to identify SNPs deviating from genome-wide population structure.

Because *BAYPASS* explicitly tests for differentiation between two groups, and additionally models read counts to account for the sampling variance inherent to Pool-Seq data, it was retained as our main outlier detection method. PCAdapt was used in parallel as a complementary exploratory tool, allowing us to verify whether similar signals emerged from unsupervised structure analyses, though its results were interpreted with caution given its limitations for Pool-Seq data.

#### BAYPASS analysis

*BAYPASS* was run with two contrasts: (1) “*in-situ*”, comparing the three Ocean pools *vs.* the three Upstream pools to test for spatial sorting in natural populations, and (2) “experimental” contrasting the pools ExpMigrant and ExpNonMigrant to test for phenotypic sorting.

We applied the procedure detailed on Figure 1. To optimize computation time, we divided the SNP dataset into 52 sub-datasets (50 random 5% samples per chromosome, plus two additional sets including missing SNPs). Each of the 52 sub-dataset was analysed with *BAYPASS* v3.0 using the option -pooldatafile and default MCMC settings. We estimated: i) the scaled covariance matrix (Ω); ii) the overall differentiation statistic *XtX*; iii) *C2* statistics (Olazcuaga et al., 2021) for both contrasts. We checked consistency across sub-datasets for Ω and hyperparameters (supplementary online Appendix 1), and results were merged after removing duplicated SNPs. The distribution of *XtX* and C2 p-values was close to uniform for high values, indicating proper calibration (supplementary online Appendix 1).

**Figure 1.**
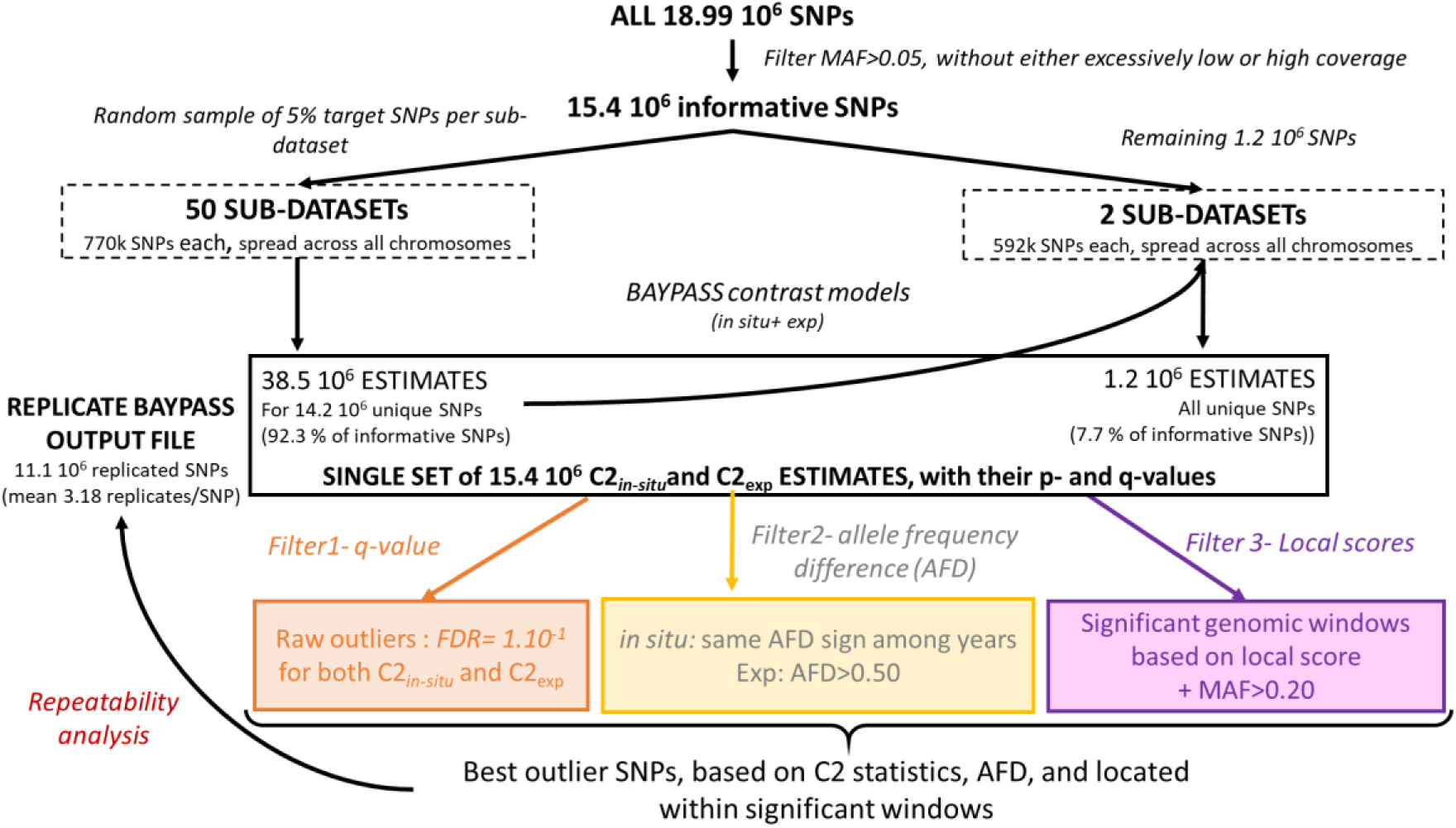
- Workflow for *BAYPASS* analyses. We split the 15.4 10^6^ SNPs into 52 subsets to run them in parallel and decrease processing elapsed time. Each subset was analysed with *BAYPASS* v3.0 using the core and contrast models (XtX and C2 statistics). Results were merged, duplicates removed, and three successive filters : (1) q-value < 1.10^-1^, consistent allele frequency differences (AFD) across years for the Ocean-Upstream contrast or AFD>0.5 (for the experimental contrast), and (3) inclusion within significant genomic windows identified by the local score method. This multi-step approach integrates SNP-level and regional signals to better capture polygenic or diffuse selection signatures.

#### Outlier filtering and local score analysis

We applied three successive filters to identify highly differentiated SNPs (Figure 1) :

1. Statistical threshold : we retained SNPs with *C2 q*-value <0.1, corresponding to a False Discovery Rate (FDR) of 10%. The R package *qvalue* was used to correct *p-*values.
2. Allele frequency differences (AFD) : for the *in-situ* contrast, we retained outliers showing consistent AFD sign across years 2019, 2020 and 2022. For the experimental contrast, only SNPs with AFD > 0.50 were kept.
3. Local score : we used the approach of Fariello et al. (2017) which accumulates modest individual signals across neighbouring SNPs to detect significant genomic windows of divergence. This method helps detect polygenic or diffuse selection signatures by identifying clusters of moderately significant markers rather than isolated outliers. We computed local scores for SNPs with MAF > 0.20, based on *C2* p-values and genomic positions, using a function adapted from Fariello et al. (2017) and available in *BAYPASS* utils (as compute.local.scores), with the parameter of the Lindley process *x_i_* =1.

SNPs that passed all three filters and remained significant in replicate analyses were retained as “best outliers”. This multi-step procedure provides a conservative test for intra-generational selection. We also applied the same procedure to identify outlier SNPs based on *XtX* statistics (with the first and third filters common for *in-situ* and experimental contrasts).

#### PCAdapt analyses

As a complementary, unsupervised analysis, we ran PCAdapt on the eight pools jointly. The dataset was also divided in 52 sub-datasets and the pcadapt function (R package *pcadapt*) was applied with *K*=1, determined by Cattell’s rule. A single z-score captures how strongly each SNP is associated with population structure, and the Mahalanobis distance (Mdist) measures each SNP deviation from the inferred population structure. We combined results from the 52 runs and applied the same three filtering steps as for *BAYPASS* (Mdist q-value < 0.1; AFD-based filtering; and local score analysis based on Mdist *p*-value with *x_i_ = 1*).

#### Comparison of F_ST_ among outliers to its empirical null distribution

To validate the outlier SNPs, we verified that their *F_ST_* values were higher than those of randomly selected SNPs. Specifically, we randomly sampled n_out_ SNPs with similar reference allele frequencies, computed *F_ST_* for each replicate, and repeated the procedure 100 times to obtain an empirical null distribution. We further extended this analysis by computing the mean *F_ST_* of the top 100, 200, … up to 1000 SNPs with the lowest *q*-values of the *C*₂ statistic, and comparing it with their empirical null distribution. *F_ST_* was estimated using the estimator for Pool-Seq data described in Hivert et al. (2018) and implemented in poolfstat (Gautier et al., 2022).

### Gene ontology analyses

We used Bedtools (2.30.0) to extract the annotations corresponding to the positions of outlier SNPs. From the nucleotide sequence codes, we searched for the corresponding gene names on NCBI and used OrthoDB v12.2 to identify orthologs genes zebrafish. Functional enrichment analyses were conducted with Enrichr and an FDR cutoff of 0.05. To assist in the synthesis and categorization of gene functions across biological processes, a large language model (ChatGPT, OpenAI) was used as a support tool to summarize and compare existing annotations. All interpretations and biological conclusions were subsequently reviewed and validated by the authors.

## Results

The whole-genome sequencing of the eight European eels’ pools revealed a total of 18.99 10^6^ SNPs after quality filtering, equating into an average of 1.94 SNPs per 100 base pairs (∼1.9% of the 979 Mb genome referenced in NCBI). The global *F_ST_*-value of 0.0086 indicates the absence of overall genetic structure among pools. In particular, *F_ST_* was ≤ 0.01 between years, and between “Ocean” and “Upstream” pools collected *in-situ* (Figure S1). Nonetheless, multi-locus *F_ST_* computed over sliding-windows of 100k SNPs show some higher values (Figure S1). Removing SNPs with either excessively low or high coverage and insufficient polymorphism reduced the genomic dataset to 15.4 10^6^ informative SNPs which were used in following analyses.

### The *in-situ* approach revealed the genomic signature of spatial sorting between Ocean *vs.* Upstream sites

Using *BAYPASS*, 26 outlier SNPs showed a significantly higher level of differentiation between Ocean and Upstream sites than expected, as indicated by the *C2_in-situ_* statistic at the 10% FDR threshold (Table S2). All of these raw outliers displayed between-sites AFD of the same sign for the three years studied. In addition, the local score approach revealed 88 significant windows of high genomic divergence extending over 10.8 kbp (Table S3, Figure S2). Combining the three filters revealed five « best outlier SNPs » within the same significant window of divergence in chromosome 1 (Figure 2). The average q-value of their *C2_in-situ_* statistic was 0.039 (Table S2).

**Figure 2.**
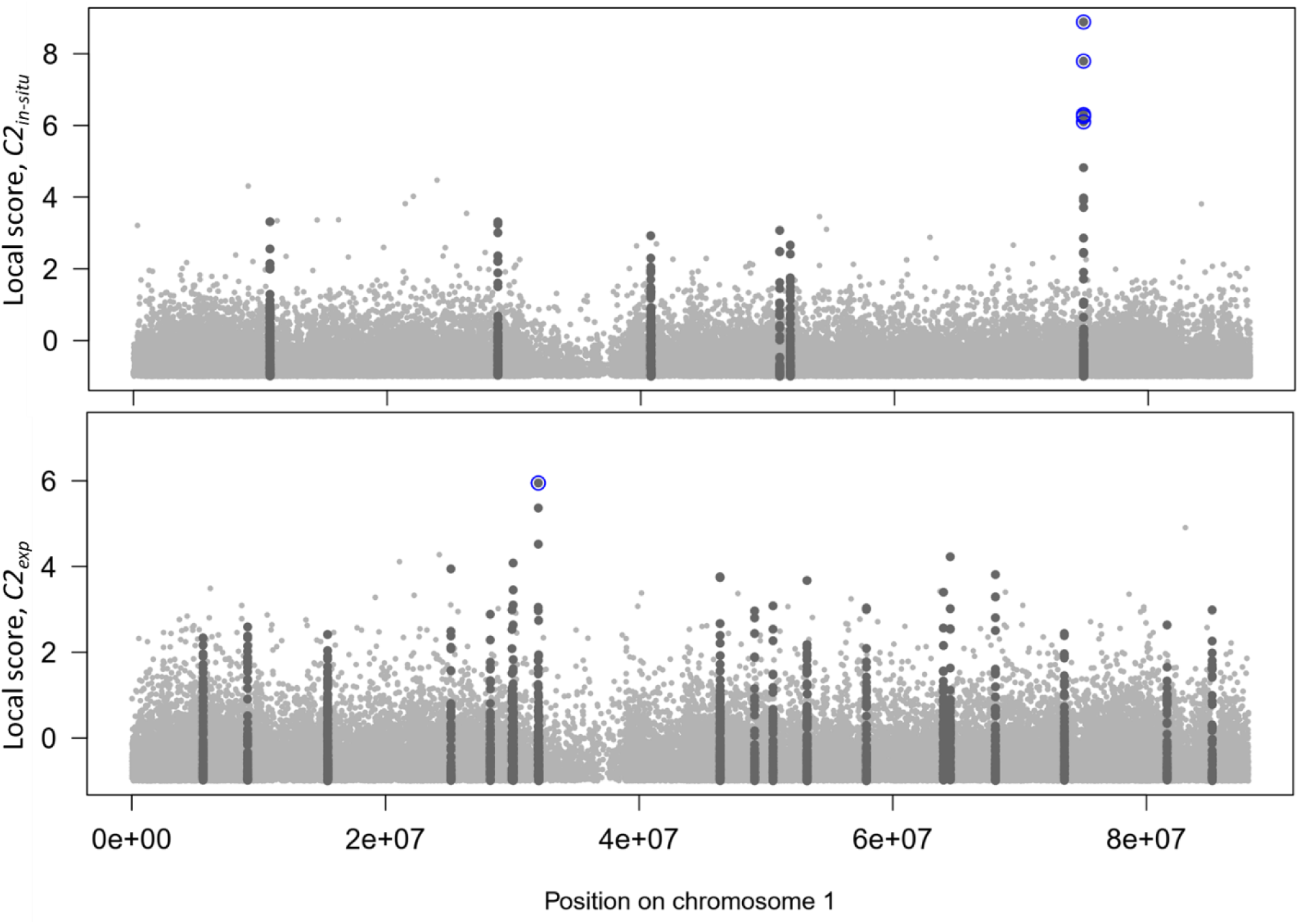
- Manhattan plots showing signatures of intragenerational selection detected with *BAYPASS* for chromosome 1 for the *in-situ* (top) and experimental (bottom) contrasts. The background local scores for all SNPs and the *C2* statistics appear in light grey points. Significant windows of high differentiation (Lindley score > 15) are marked by dark grey points. Blue circles highlight the best outliers SNPs, which passed the three filters (q-value, AFD, local score) described in Figure 1.

These five best outlier SNPs were confirmed when using *BAYPASS* with the *XtX* statistics and similar filtering criteria (Table S2, S3, Figure S3). Using the complementary approach *PCAdapt*, we identified 2110 outlier SNPs (Table S2). Among them, 592 displayed between-sites AFD of the same sign for the three years studied. Moreover, the local score approach detected 1069 significant windows of high genomic divergence extending over 9.9 kb (Table S3, Figure S2). Combining the three filters revealed 239 « best outlier SNPs ». One of them was common with those identified by *BAYPASS* (Figure S3). Overall, outlier detection based on *BAYPASS* and the *C2_in-situ_* statistic appears to be the most conservative approach, as it identifies a very limited number of SNPs that are independently supported by the XtX statistic, whereas the alternative methods PCAdapt detects a much larger set of candidates that likely include a substantial proportion of false positives. For this reason, we focused subsequent analyses on the five best outlier SNPs consistently detected by *BAYPASS*.

The five best outlier SNPs showed contrasted frequencies in Ocean *vs.* Upstream sites (Figure 3). On average across SNPs and years, allele frequencies increased from 0.577 in Ocean pools to 0.96 in Upstream pools. This variation from intermediate to extreme frequency is consistent with our hypothesis that Ocean sites host a mixture of migrant and non-migrant glass eels, while Upstream sites host mainly migrant glass eels. The synchronous variation across neighbor SNPs is likely due to a combination of spatial sorting and linkage disequilibrium.

**Figure 3.**
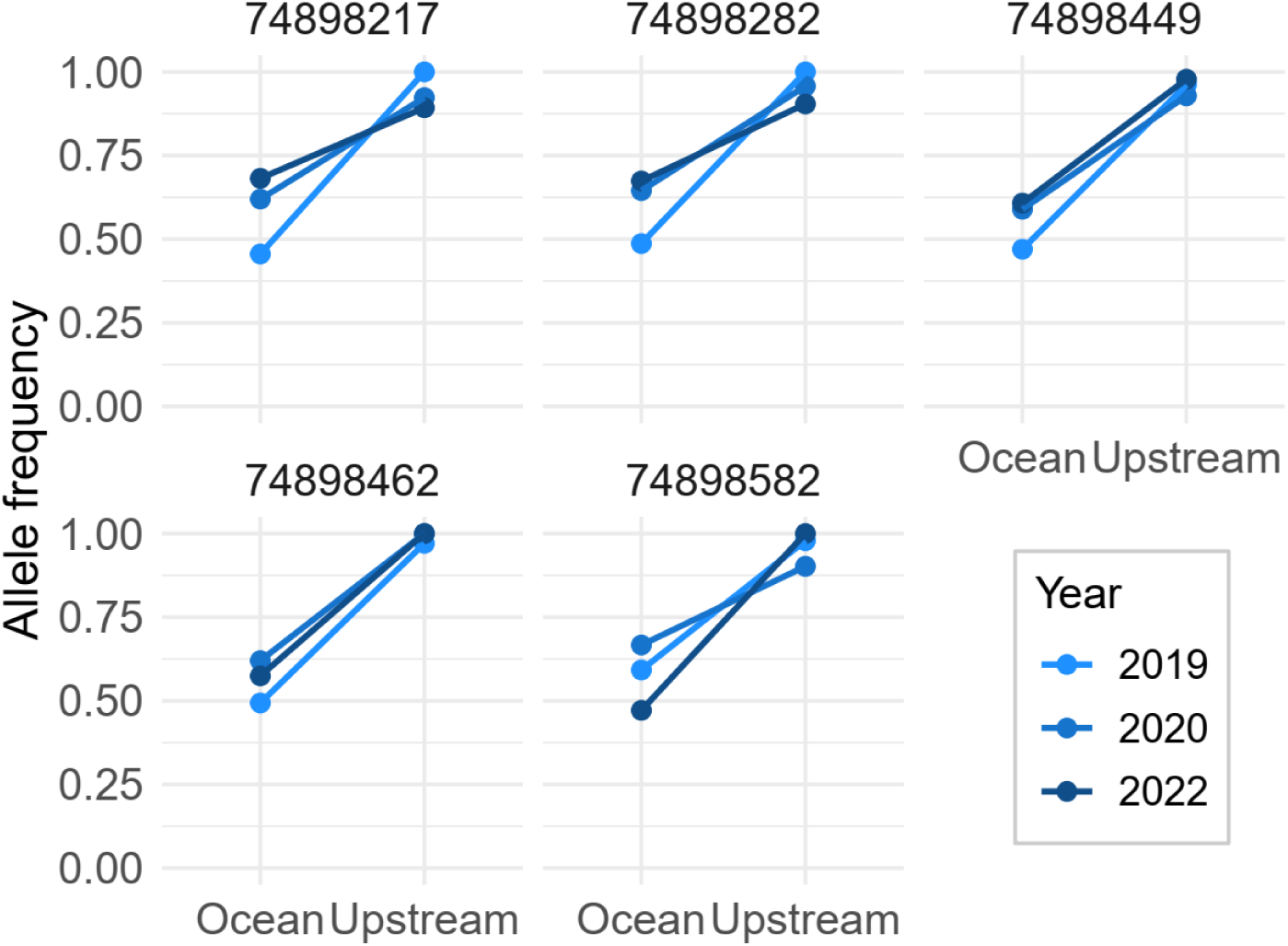
- Variation of allele frequencies at the five best *in-situ* SNPs (identified by their position on chromosome 1). Allele frequencies in each pool (Ocean vs Upstream) are shown for the five outlier SNPs. Points and lines connect the frequencies across the three sampling years. For simplicity, this representation assumes that the proportion of reads carrying a given allele provides an unbiased estimate of allele frequency.

The mean between-site *F_ST_* at the five best outlier SNPs was 0.21, significantly higher than the empirical null distribution obtained after controlling for allele-frequency effects through random SNP sampling (Figure 4). Moreover, the signal of high differentiation was not restricted to these five best outlier SNPs: it persisted when considering the top 100, 200, and up to 1000 SNPs with the lowest *q*-values of the *C2_in-situ_* statistic. Mean *F_ST_* steeply declined to 0.11 for the top 100 SNPs, and then more gradually (e.g. *F_ST_* =0.079 for the top 1000 SNPs).

**Figure 4.**
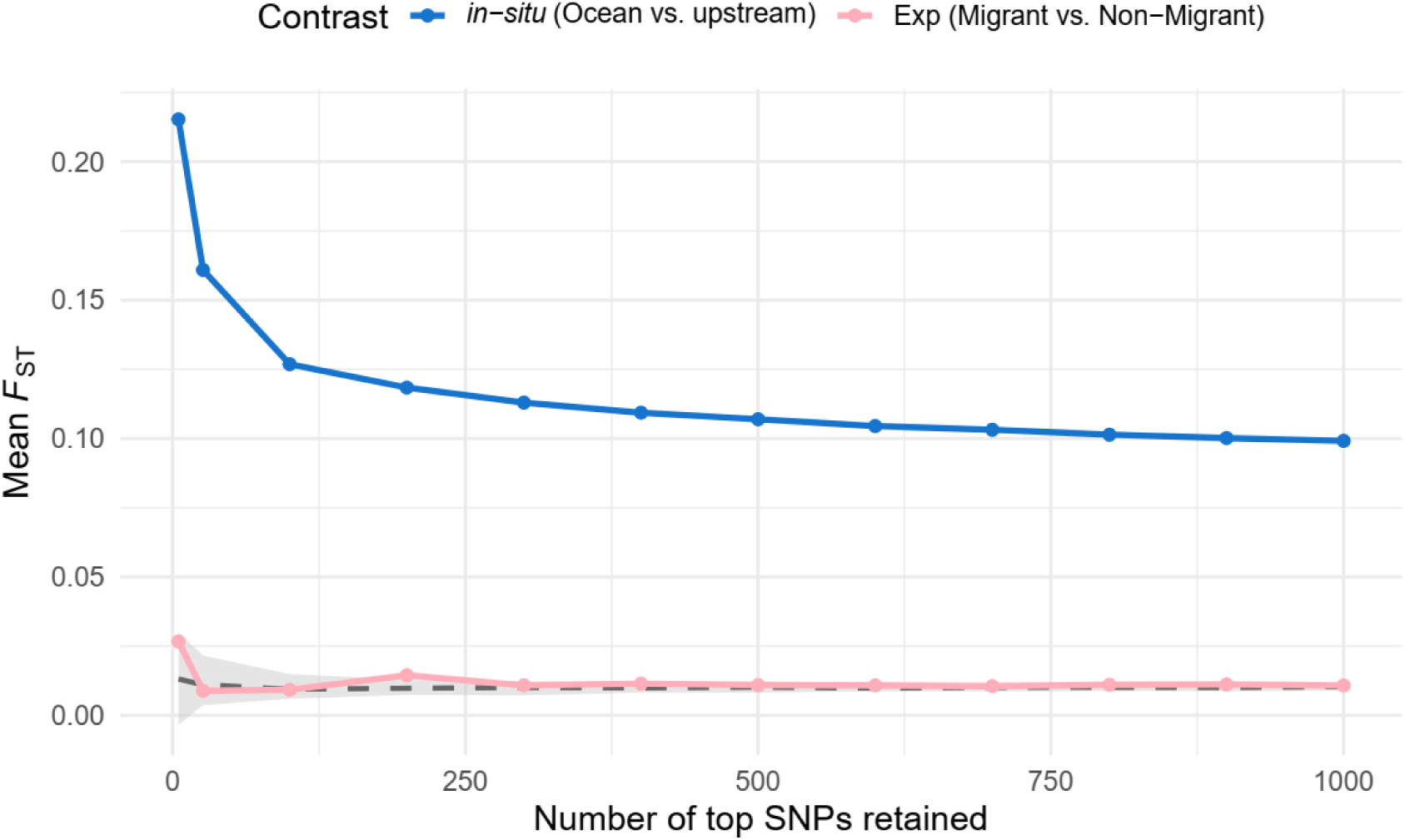
- Variation of mean *F_ST_* values computed for different sets of top *in-situ* outlier SNPs. The blue lines corresponds to average *F_ST_* between Ocean and Upstream sites (*in-situ*), and pink lines to *F_ST_* between experimental Migrant and Non-migrant pools. The grey envelope represents the neutral expectation for *F_ST_* between Ocean and Upstream sites; the dashed line indicates its median value. We considered successive sets of SNPs identified from the *in-situ* contrast: the 5 “best” outliers, and then the top 26, 100, 200, …, 1000 SNPs with the lowest q-value. smuratorioTo build the neutral envelope, we randomly sampled, for each SNP set size, an equivalent number of loci with similar allele frequencies, and computed *F_ST_*. This was done 100 times to derive the 2.5–97.5% quantile range (grey area). See also Figure S4 (larger number of top SNPs).

### The experimental approach revealed genomic signature of phenotypic sorting between migrant and non-migrant eels

Using *BAYPASS*, 1090 outlier SNPs showed significant differentiation between the ExpMigrant and ExpNonMigrant pools as indicated by the *C2_exp_* statistic at the 10% FDR threshold (Table S2). Among them, 200 showed AFD>0.5 between pools. Finally, 219 windows of high differentiation spanning over 10.8 kbp (median) were found using the local score approach. By combining the three filters, 28 experimental « best outlier SNPs » were identified in 13 chromosomes (Figures 1 and S5). Relaxing the second filter to AFD>0.25 resulted in 49 best outlier SNPs.

Using *BAYPASS* with the *XtX* statistics and similar filtering criteria, only five best outlier SNPs were retained, two of which overlapped with the 28 SNPs identified using the *C2_exp_* statistic (Table S2, S3, Figure S3). Using the complementary approach *PCAdapt*, we identified 19 « best outlier SNPs », none of which overlapped with those detected by *BAYPASS* using the *C2_exp_* statistic (Figure S3). Overall, these discrepancies likely reflect limited statistical power associated with the use of only two experimental pools. For consistency with the in situ analytical strategy, subsequent analyses therefore focused on the 28 best outlier SNPs identified using *BAYPASS* and the *C2_exp_* statistic.

These 28 best outlier SNPs showed contrasted frequencies in the ExpMigrant *vs.* ExpNonMigrant pool (Figure S6), with a mean *F_ST_* of 0.48, significantly higher than the empirical null expectation (Figure S7). As for in-situ outliers, the signal of high differentiation persisted when considering the top 100, 200, and up to 1000 SNPs with the lowest *q*-values of the *C2_exp_* statistic (Figure S7).

### Functional interpretation of intragenerational selection signatures and overlap between the *in-situ* vs experimental approaches

The five best outlier SNPs identified *in-situ* were all located within intronic regions of the same gene, whose zebrafish orthologs (e.g, gpb2) encode a guanylate-binding protein involved in immune system process (Table S4A).

In contrast, the 28 best outliers SNPs identified in experimental conditions were distributed accross 23 distinct genomic regions. One SNP corresponded to a synonymous variant, 11 were located in introns while the remaining SNPs intergenic (3 SNPs), or located upstream or downstream of the genes (13 SNPs), possibly affecting regulatory regions. Over the 28 genes containing or located close to these SNPs, 22 had identifiable zebrafish orthologs (Table S4B). Based on functional annotations, these genes could be broadly grouped into three main functional categories: (1) neuronal development (*filip1l, pcdh17, sema7a, sv2bb, isl2b, plppr4a, FAM126Bl*); (2) cell migration, cytoskeleton organization and tissue remodeling (*filip1l, sema7a, pcdh17, crocc, hspg2, ptpn11a*); (3) Transcription/RNA processing and DNA reparation (*tdrd3, ctdp1, cenpo, parp*) . However, gene ontology enrichment analyses did not yield statistically significant terms after multiple testing correction, likely due to the limited size and functional heterogeneity of the gene set. In addition, two genes displayed annotations particularly relevant in our context: *crat*, involved in lipid and energy metabolism, and *dbp*, a transcription factor implicated in the regulation of circadian rhythms.

No best outlier SNPs or associated genes were shared between the *in-situ* and experimental contrasts (Figure S8). When extending the comparison to the top 100 outlier SNPs identified by each approach, only one genomic region (intergenic) was shared between contrasts (Figure S8). The two genes flanking this region have zebrafish orthologs: ikzf2, involved in metal ion binding, and vwc2l, implicated in nervous system development.

## Discussion

By combining *in-situ* and experimental approaches, this study provides new insights into the genetic determinants of selection on estuarine migration propensity in European glass eels viewed from a whole-genome perspective. The *in-situ* approach identified five *F_ST_*-outlier SNPs in one single gene that consistently showed high differentiation between the estuary mouth and a freshwater site Upstream over three years (*i.e.,* spatial sorting). Additionally, 28 *F_ST_*-outlier SNPs distributed across 28 genes were significantly differentiated between migrant and non-migrant glass eels in experimental conditions (*i.e.,* phenotypic sorting). The absence of overlap between the *in-situ* and experimental approaches highlights several possible challenges for further research into the physiological mechanisms underlying migration in diadromous fish.

### Intragenerational selection on the propensity to migrate along the estuary in European eel

In accordance with panmixia, we found a high genetic diversity in European eel, with a total number of 18.99 × 10^6^ SNPs for a genome size of 0.979 Gb, equating into an average of 1.94 SNPs per 100 base pairs. Together with the absence of genetic structure among pools sampled in different years, these results confirm the significant role of panmixia in shaping genetic diversity and structure in this species, a topic of extensive debate (Schmidt, 1923; Avise et al., 1986; Wirth & Bernatchez, 2001; Dannewitz et al., 2005), now well established (Gagnaire et al., 2012; Pujolar et al., 2014; Pavey et al., 2015; Babin et al., 2017; Enbody et al., 2021).

This exploratory study provides evidence of genotype-phenotype and genotype-environment associations, suggesting spatially variable selection acting on estuarine migration propensity in European glass eels. Our hypothesis was that migration events can act as strong selective filters by spatially sorting individuals according to their migration ability, behaviour, and associated functional traits. We define here the propensity of glass eels to migrate within estuary as being influenced by four underlying traits category: (i) swimming ability including factors like energetic status (Bureau Du Colombier et al., 2007; Edeline, 2007); (ii) ability to synchronize swimming activity with the tide, which involves sensory perception and circatidal rhythms (Creutzberg, 1959; McCleave & Kleckner, 1982; Wippelhauser & McCleave, 1987; Bolliet et al., 2007; Bolliet & Labonne, 2008), (iii) ability to detect and respond to varying environment conditions along the estuary such as changes in temperature, hydraulic conditions, salinity, microbial community or pollution (Wilson et al., 2004; Sasai et al., 2007) and (iv) decision-making traits that affect their migratory behaviour such as habitat choice eventually triggered by population density (Pavey et al., 2015).

The five outliers SNPs showing significant differentiation between the mouth of the Adour estuary and the Upstream site located 40km away, as observed in the *in-situ* approach, could be attributed to genomic signatures of spatial sorting acting on any one of these four traits, or on their combination. The 28 outliers SNPs showing significant differentiation between migrant and non-migrant glass eels in the experimental approach suggest significant genotype-phenotype relationship and could be specifically related to a genetic determinism of glass eels’ ability to swim and to synchronize their activity with the change in water current direction mimicking changing tides (traits *i* and *ii* mentioned above). However we cannot rule out that traits related to manipulation and perception of experimental environment are also involved in this experimental contrast.

Our *in-situ* and experimental contrasts are thus highly complementary, and comparison of their associated sets of outliers SNPs and genes brings several insights. From a quantitative perspective, the fact that we detected a single gene under spatially variable selection in the *in-situ* approach, compared to 28 genes associated with migration propensity in the experimental approach, may have several explanations. First, the *in-situ* contrast, based on three pairs of Ocean versus Upstream sites replicated across years, is statistically more stringent than the experimental contrast based on a single pair. In addition, it searches for signatures of intragenerational selection that are consistent overs years, independently of interannual variability in environmental conditions (e.g. related to temperature or flow). Second, the experimental contrast, analogous to a Genome-Wide Association Study, focusses on extremes values of a restricted number of traits (swimming ability and synchronization of activity) in constant conditions using only the change in water current direction as tidal *zeitgeber* to synchronise swimming activity. In contrast, the *in-situ* contrast may capture a more integrated response involving a larger number of traits, including response traits to varying environment conditions along the estuary. When the number of causative loci is high, as expected for complex, integrated traits, individual loci become increasingly difficult to detect because the strength of selection acting on each of them is reduced, leading to only small allele frequency shifts. The progressive decline of mean *F*_ST_ when expanding the set of candidate SNPs (Figure 4) suggests a diffuse but consistent genomic signal of spatial sorting, compatible with weak intragenerational selection acting on multiple loci, partly through linkage disequilibrium. Instead, it has been proposed that the response of integrated traits to selection derives mostly from allelic co-variances among causative loci (Mckay, Latta & Mckay, 2002; Kremer & Corre, 2011). The importance and role of these allelic co-variances could not be addressed in this exploratory Pool-Seq study and should be investigated further in future population genomic analyses.

The absence of overlap between the *in-situ* and experimental set of outlier genes could also be explained by several other factors. Before turning to functional interpretation, developed in the next paragraph, we note that limited power in the *in-situ* approach may have prevented the detection of some loci common to both contrasts (see above). In addition, because the experimental approach relies on glass eels captured at the mouth of the Adour estuary, and then sorted in freshwater conditions, genotype–phenotype interactions cannot be excluded, whereby certain genotypes may express different migration abilities or behaviours depending on environmental context. To address this issue, future studies could, for instance, experimentally sort glass eels in saltwater, or glass eels collected at the upstream site, which would help refine our understanding of the relationship between phenotypic and spatial sorting.

This study also shares the classical limitations of population genomic approaches, in which selection is inferred from allele frequency changes rather than directly measured through differential fitness across environments or phenotypes. Moreover, although this was precisely our objective, it is important to note that we focused on allelic frequency variation occurring over a very short time window and at an early life stage of the eel life cycle, bearing in mind that not all our migrant or non-migrant individuals will eventually contribute to reproduction. Beyond the few loci showing signatures of spatially variable selection, most of the genome remains weakly differentiated between the mouth of the Adour estuary and the upstream site, indicating that substantial genetic variation remains available all along the estuary for potential responses to selection at later life stages. Finally, an important unresolved issue concerns condition-dependent filtering through mortality in each environment. Here, we assume that genetic differentiation primarily reflects genotype-dependent physiological ability to migrate and/or genotype-dependent habitat choice, two processes that are difficult to disentangle without targeted experimental approaches beyond the scope of this study (see also Pavey et al., 2015). However, differential mortality or settlement along the estuarine gradient may also contribute, a demographic process that remains largely undocumented for glass eels.

### Tentative interpretation of the biological processes involved in migration propensity in eels

The functional annotation of outlier genes revealed by *in-situ* and experimental approaches can provide useful insights into the genomic pathways potentially associated with migration propensity in eels. However, these interpretations must be treated with caution, given the absence of direct functional validation of these genes in *Anguilla anguilla*, the absence of overlap, and thus of reciprocal support, between the *in-situ* and experimental sets of outlier genes, and the well-known risk of over-interpretation or “storytelling” in population genomics (Pavlidis et al., 2012).

Focusing first on the 22 functionally annotated genes associated with contrasted phenotypes in the experimental approach, two functional categories appear particularly consistent with the phenotypic sorting on swimming ability and synchronization of activity. First, seven genes are involved in biological processes related to neuronal development. These genes may contribute to the establishment and modulation of neural circuits underlying motor coordination, sensory integration, and behavioural responses required during active migration. Second, six genes are involved in cell migration, cytoskeleton organization, and tissue remodelling. Although these genes act at different molecular and cellular levels, they converge on processes controlling cellular mechanics, adhesion, and signal integration, all of which are essential for effective locomotion and sustained swimming performance. Taken together with the two genes involved in lipid and energy metabolism (crat), as well as in the regulation of circadian rhythms (dpb), this set of candidate genes draws a coherent picture of biological processes associated with migratory locomotion. These processes span multiple levels of biological organization, from energetic supply and metabolic regulation to neuronal activity and temporal coordination, all of which are indispensable for orientation and movement in a fluctuating estuarine environment.

By contrast, the single outlier gene identified in the *in-situ* approach was involved in antiviral immune response. This result is consistent with the high physiological stress experienced by glass eels during estuarine migration, particularly the osmoregulatory cost associated with the transition from saline to freshwater environments. Such stress is likely to increase susceptibility to pathogens and disease. Although only a limited number of viruses have so far been documented in *A anguilla* (McConville et al., 2018), genetic variation in immune-related genes may nevertheless influence individual survival by modulating vulnerability to disease-related mortality during this critical life stage. By comparison with the functional annotations of outlier genes of experimental approach, those of the in-situ approach suggests that eel ability to swim or to synchronise with tides may not be the limiting factor for colonising upstream of the estuary.

The prominence of genes related to central nervous system development in the phenotypic sorting experiment of glass eels may reflect our focus on this specific life stage. In the American eel, Pavey et al. (2015) also identified genes linked to development, but more related to cardiac and respiratory system development, and limb bud formation. They also detected a function linked to olfaction, suggesting sensory perception differences. Similarly, our findings, which include brain or forebrain development could also imply variations in sensory perception.

### Synthesis, and implication for management and beyond

Consistent with the panmictic nature of European eel populations, genome-wide differentiation between Ocean and Upstream sites remains extremely low. This indicates that, beyond a limited number of candidate SNPs, most of the genome is effectively homogenized by gene flow, and that spatially variable selection within the estuary, if present, leaves only weak and localized genomic signatures. At first glance, this weak genomic signature of spatially variable selection detected *in-situ* may appear inconsistent with the stronger genotype–phenotype associations revealed under experimental conditions. However, these two results are not necessarily contradictory and may instead reflect different components of the migratory process. The experimental approach primarily captures intrinsic variation in migratory-related traits, such as swimming performance and synchronization with water currents, whereas the *in-situ* approach reflects the realized outcome of migration within a complex and heterogeneous estuarine environment. Importantly, the ability to migrate may not be sufficient, nor even necessary, to determine upstream movement within the estuary. Some glass eels that are physiologically capable of migrating may not colonise upstream areas if other conditions are not met, such as appropriate hormonal or energetic status (Edeline et al., 2004; Bureau Du Colombier et al., 2007). Conversely, some glass eels with a lower intrinsic migratory capacity may still reach upstream habitats under favourable local environmental conditions or due to phenotypic plasticity (e.g., health status). Such genotype-by-environment interactions may dilute the genomic signature of spatial sorting, without questioning the existence of a genetic basis for migratory traits.

There is considerable conservation concern for the European eel, which has led to costly annual restocking operations both upstream and downstream within river basins, and sometimes among basins. These restocking operations generally target sites considered favourable for growth and characterized by low eel densities. However, eel density and environmental conditions vary strongly along the longitudinal estuarine gradient, particularly with respect to salinity, tidal influence, and hydrodynamics. Our results, highlighting a genetic basis for migration propensity and a moderate but detectable effect of intragenerational spatially variable selection during estuarine migration, suggest that not all glass eels have the same ability to migrate or to cope with environmental conditions encountered upstream or downstream in the estuary. More broadly, these findings are consistent with previous population genomic studies showing intragenerational adaptive differentiation in European eel at larger geographical scales, notably among river basins (Gagnaire et al., 2012; Pujolar et al., 2014). While these earlier studies emphasized the role of climatic factors, particularly temperature, in shaping large-scale latitudinal sorting of eels along the coast, our study focuses on intrinsic traits such as migration ability and habitat choice driving longitudinal spatial sorting within estuaries. Taken together, these population genomic insights call for caution when implementing restocking operations, as they implicitly assume functional equivalence among stocked individuals across contrasted environments.

Overall, this exploratory study of the genomic basis of migration propensity in eel highlights several avenues for further investigation. Increasing sampling density along the Adour estuary could help clarifying whether allelic frequencies shift uniformly along the gradient or if specific stages of differentiation, potentially influenced by salinity or tidal dynamics, exist. Experimentally sorting glass eels collected at upstream sites could further elucidate genotype-by-environment interactions affecting migratory behaviour. Finally, extending this study across multiple estuaries along a latitudinal gradient could help assess the generality and robustness of the intragenerational selection patterns observed in the Adour estuary, and clarify the conditions under which such processes can be detected.

## Acknowledgements

We are grateful to the GeT-PlaGe platform (INRAE, Toulouse, France), member of the GeT Genotoul platform, for sequencing support and the Sigenae bioinformatics platform (INRAE, Toulouse, France), member of the GeT Genotoul platform, for providing computing and storage resource. We are grateful to the genotoul bioinformatics platform Toulouse Occitanie (Bioinfo Genotoul, https://doi.org/10.15454/1.5572369328961167E12) for providing computing and storage resources. We thank our colleagues J Labonne, F Daverat and C. Bouchard for useful discussions and comments on a previous version of the manuscript.

## Funding

S. Sebihi’s PhD thesis was funded by the University of Pau and Pays de l’Adour (UPPA) through the Transborder PhD programme, in partnership with the University of the Basque Country (UPV/EHU). Whole-genome PoolSeq sequencing was funded by INRAE through the internal funding scheme “Pari scientifique 2023”.

## Conflict of interest disclosure

The authors declare that they comply with the PCI rule of having no financial conflicts of interest in relation to the content of the article. SOM is recommenders for two Peer Communities (PCI Evol Biol; PCI Ecology).

## Data, scripts, code, and supplementary information availability

Raw sequencing data have been deposited in the European Nucleotide Archive (ENA) under accession number PRJEB106189.

Scripts, codes and Supplementary online Appendix 1 are available online: https://doi.org/10.57745/TDULXV (Oddou-Muratorio et al., 2026).

Supplementary Figures and Tables for this article are available at the end of this file.

## CRediT authorship contribution statement

**Stellia Sebihi**: Conceptualization, Methodology, Software, Formal Analysis, Investigation, Data Curation, Visualization, Writing – Original Draft. **Aurélie Manicki**: Methodology, Software, Resources, Writing – Review & Editing. **Christophe Klopp**: Methodology, Software, Resources, Data Curation, Writing – Review & Editing. **Mathieu Gautier**: Methodology, Software, Resources, Writing – Review & Editing. **Pascale Coste** : Methodology, Resources, Writing – Review & Editing. **Emmanuel Huchet** : Methodology, Resources, Writing – Review & Editing. **Maren Ortiz-Zarragoitia** : Writing – Review & Editing, Supervision. **Valerie Bolliet**: Conceptualization, Investigation, Data Curation, Supervision, Funding Acquisition, Writing – Review & Editing. **Sylvie Oddou-Muratorio**: Conceptualization, Methodology, Software, Formal Analysis, Data Curation, Visualization, Supervision, Funding Acquisition, Writing – Review & Editing.

## Supplemental information for the manuscript “Weak but consistent genomic signals of intragenerational selection during estuarine migration in the European eel (*Anguilla anguilla*)”

**Table S1:**
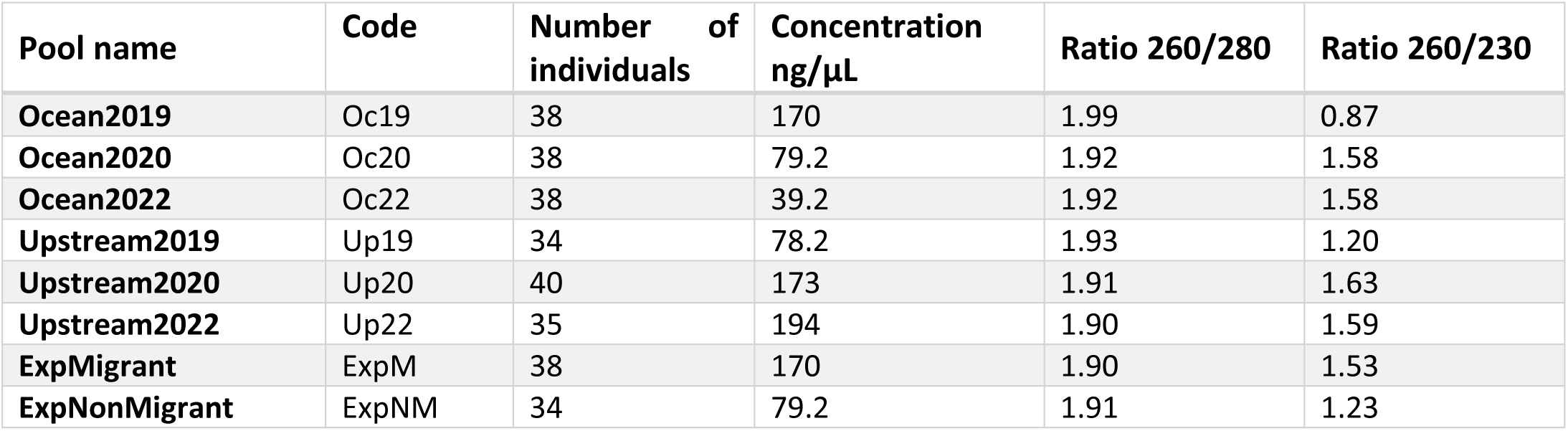
Code, DNA quantity and quality of each pool. Concentration and ratios were measured with Qbit 2.0 fluorometer (Thermo fisher Scientific) and nanodrop. The 260/280 and 260/230 ratios relate to the purity of DNA samples.

| **Pool name** | **Code** | **Number of individuals** | **Concentration ng/μL** | **Ratio 260/280** | **Ratio 260/230** |
| --- | --- | --- | --- | --- | --- |
| **Ocean2019** | Oc19 | 38 | 170 | 1.99 | 0.87 |
| **Ocean2020** | Oc20 | 38 | 79.2 | 1.92 | 1.58 |
| **Ocean2022** | Oc22 | 38 | 39.2 | 1.92 | 1.58 |
| **Upstream2019** | Up19 | 34 | 78.2 | 1.93 | 1.20 |
| **Upstream2020** | Up20 | 40 | 173 | 1.91 | 1.63 |
| **Upstream2022** | Up22 | 35 | 194 | 1.90 | 1.59 |
| **ExpMigrant** | ExpM | 38 | 170 | 1.90 | 1.53 |
| **ExpNonMigrant** | ExpNM | 34 | 79.2 | 1.91 | 1.23 |

**Figure S1:**
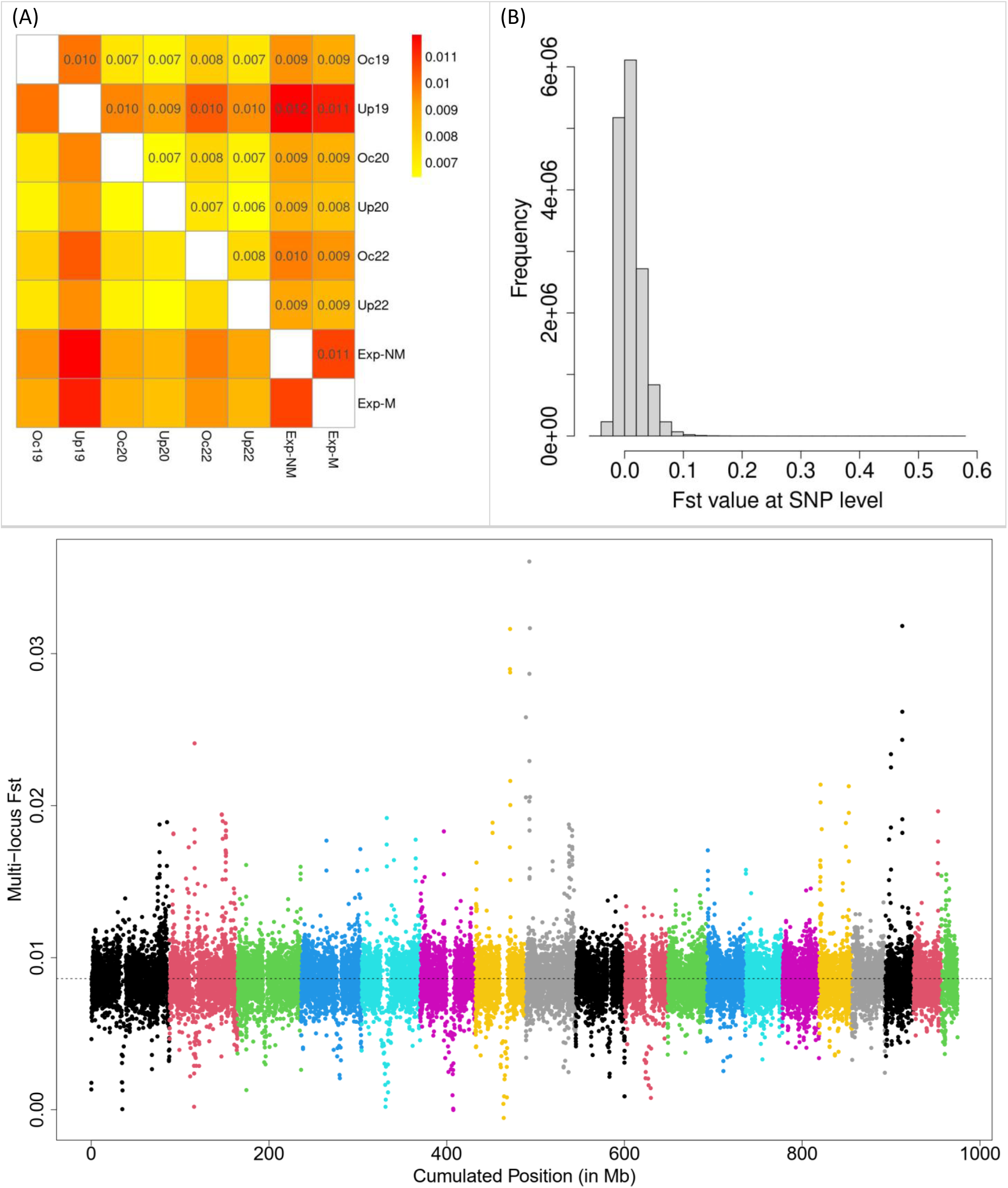
Global genomic differentiation between studied pools as measured by *F_ST_*. (A) Whole-genome pairwise *F_ST_* among the eight studied pools “(see Table S1 for code). (B) distribution of individual SNP- *F_ST_* values and (C) Manhattan plot of the multi-locus *F_ST_* computed over sliding-windows of 100k SNPs. The dashed line indicates the estimated overall genome-wide *F_ST_*. The 19 chromosomes are represented by alternate colours.

**Table S2:**
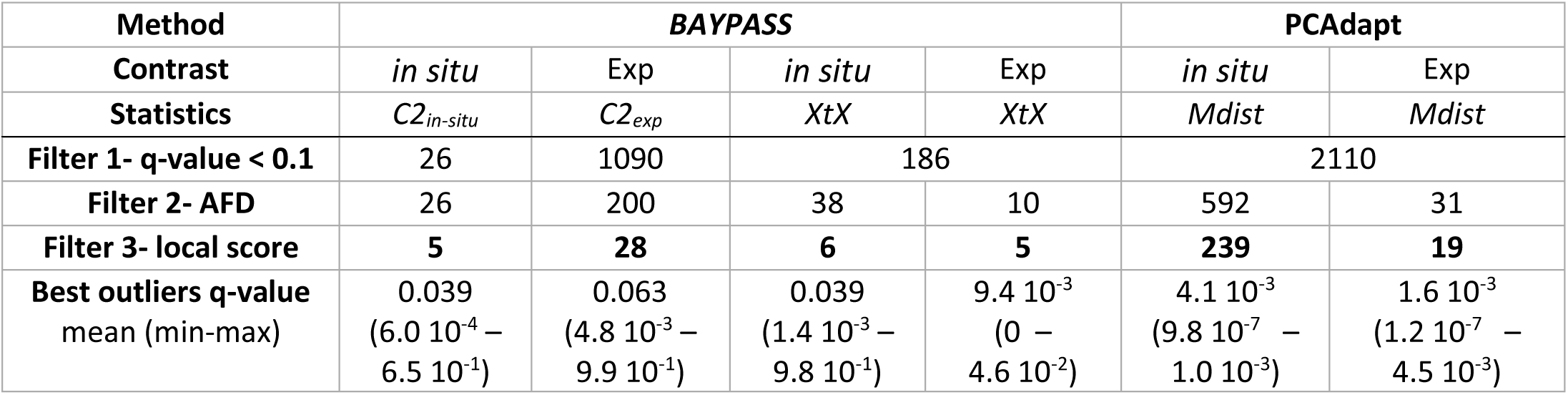
Number of outlier SNPs detected with *BAYPASS* and *PCAdapt*. Three successive filters were applied (see Figure 1 and methods for details): (1) significant outlier SNPs were identified based on a 0.1 q-value threshold (i.e., False Discovery Rate, FDR, of 1%); (2) these outlier SNPs were filtered on allele frequency difference (AFD) between contrasted pools; and (3) finally, we retained as best outliers only outlier SNPs located within genomic windows of significant differentiation, defined using the local score approach (Table S3 and Figure S2). This table presents the number of outlier SNP retained at each filtering steps, allowing comparison between *BAYPASS* (C₂ and XtX statistics) and PCAdapt (Mahalanobis distance, Mdist) results. Note that XtX- and Mdist-based results are the most comparable, as both statistics test for overall differentiation between pools (hence the single column for Filter 1 results). The last line summarizes the distribution of q-values of the best outliers SNPs (mean, minimal and maximal value in brackets). **Overall, *PCAdapt* detected a higher number of outlier SNPs than *BAYPASS* for the same FDR threshold, and this difference persisted after the three filtering steps, reflecting the lower stringency of the local score filter applied to PCAdapt results (see Table S3)**.

| Method | *BAYPASS* |  |  |  | *PCAdapt* |  |
| --- | --- | --- | --- | --- | --- | --- |
| Contrast | *in situ* | Exp | *in situ* | Exp | *in situ* | Exp |
| Statistics | $C_{2in-situ}$ | $C_{2exp}$ | $XtX$ | $XtX$ | $Mdist$ | $Mdist$ |
| **Filter 1- q-value < 0.1** | 26 | 1090 | 186 |  | 2110 |  |
| **Filter 2- AFD** | 26 | 200 | 38 | 10 | 592 | 31 |
| **Filter 3- local score** | **5** | **28** | **6** | **5** | **239** | **19** |
| **Best outliers q-value** | 0.039 | 0.063 | 0.039 | $9.4 \cdot 10^{-3}$ | $4.1 \cdot 10^{-3}$ | $1.6 \cdot 10^{-3}$ |
| mean (min-max) | $(6.0 \cdot 10^{-4} - 6.5 \cdot 10^{-1})$ | $(4.8 \cdot 10^{-3} - 9.9 \cdot 10^{-1})$ | $(1.4 \cdot 10^{-3} - 9.8 \cdot 10^{-1})$ | $(0 - 4.6 \cdot 10^{-2})$ | $(9.8 \cdot 10^{-7} - 1.0 \cdot 10^{-3})$ | $(1.2 \cdot 10^{-7} - 4.5 \cdot 10^{-3})$ |

**Table S3:** Characteristics of genomic windows of high differentiation identified by local score analyses. For each method and test statistics, the local score approach identified the total number of significant windows (# win), their size (median [min – max]), the total number of SNPs included, and the mean Lindley score and mean –log₁₀(p-value) of SNPs within these windows. We chose a Lindley score threshold of 15, as determined through graphical sensitivity analyses (see Fig. S3). **Overall, local score analyses revealed a higher number and larger size of significant windows for *PCAdapt* as compared to *BAYPASS* results.**

| Method | *BAYPASS* |  |  | *PCAdapt* |
| --- | --- | --- | --- | --- |
| Contrast / Global test | *in situ* | Exp | Global | Global |
| Statistics used | $C_{2in-situ}$ | $C_{2exp}$ | $XtX$ | $Mdist$ |
| **# windows** | 88 | 219 | 34 | 1069 |
| **Window size (kbp)** | 10.8 | 10.8 | 4.9 | 9.9 |
| **[min-max]** | [935 bp -451 kbp] | [931 bp -266 kbp] | [1015 bp -50 kbp] | [895 bp -1 Mbp] |
| **# SNPs** | 7 880 | 17 376 | 2 200 | 111 512 |
| **Mean Lindley score** | 21.2 | 20.8 | 21.0 | 25.1 |
| **Mean $-\log_{10}(p\text{-value})$** | 4.0 | 4.3 | 5.0 | 5.3 |

**Figure S2:**
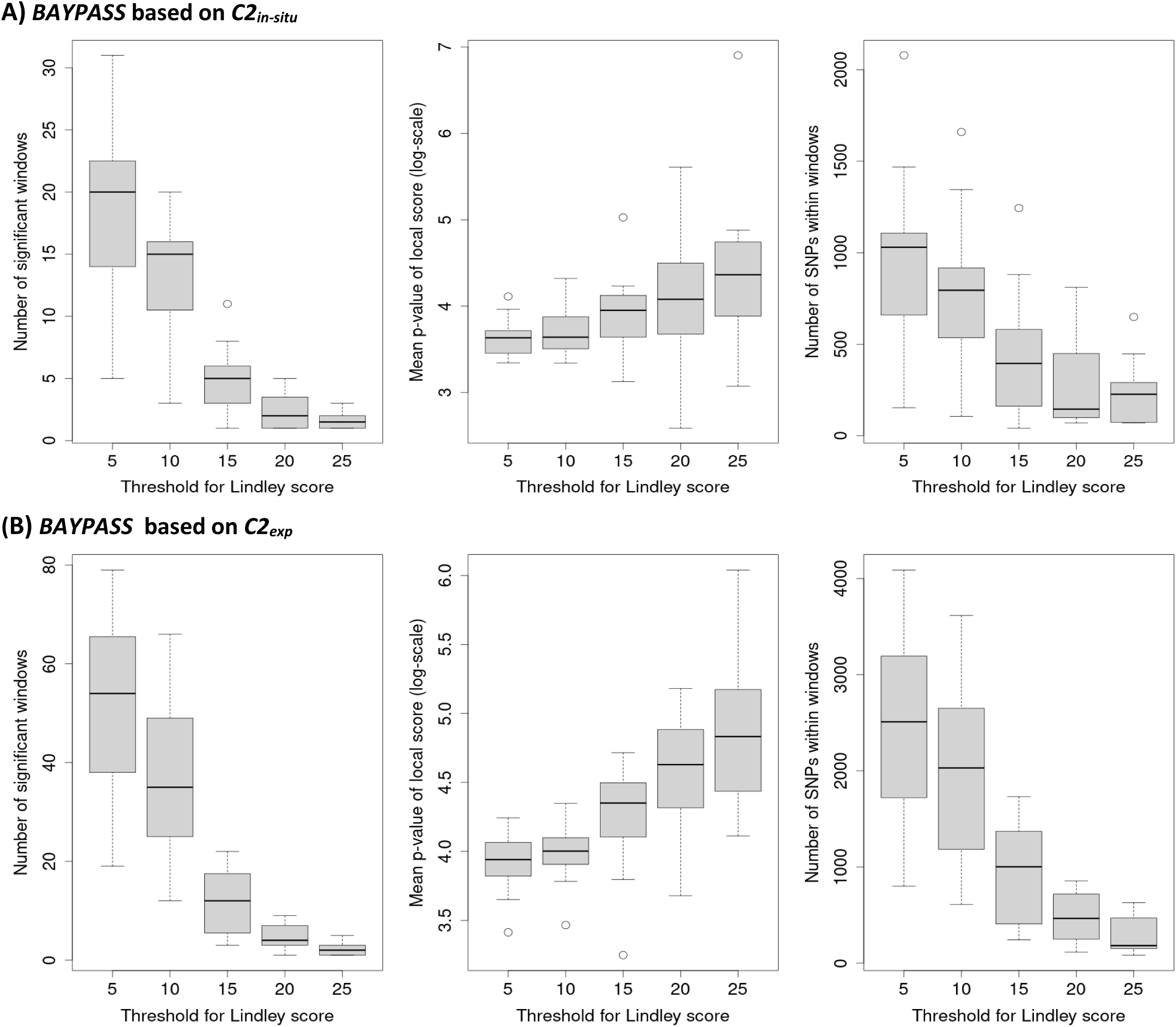
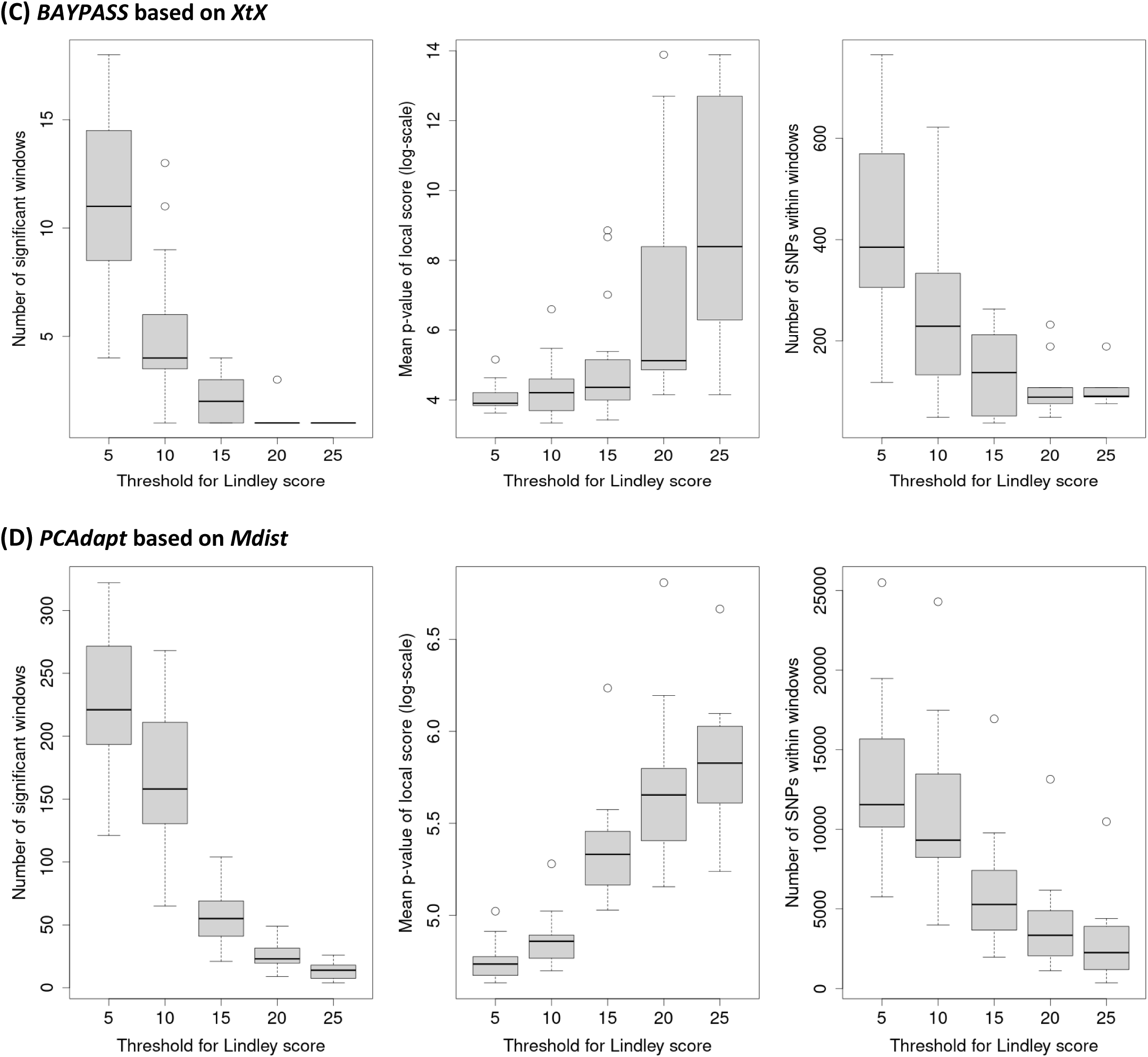
Sensitivity analyses of local score approach used in *BAYPASS* analyses. Graphical sensitivity analyses were conducted to determine the threshold value of the Lindley score used to define significant genomic windows of high differentiation. The boxplots show how three summary metrics vary with the Lindley score threshold: (i) the number of windows of high differentiation (left), (ii) the mean –log₁₀(p-value) of SNPs within these windows (middle), and (iii) the total number of SNPs within these windows (right), for each method: (A) *BAYPASS* based on *C2_in-situ;_*; (B) *BAYPASS* based on *C2_exp_*; (C) *BAYPASS* based on *XtX*; (D) *PCAdapt* based on *Mdist* **A marked decrease in the number and size of significant windows was observed between Lindley score thresholds of 10 and 15 for panels A, B, and D. We therefore retained a conservative threshold of 15 for outlier detection.**

**Figure S3:**
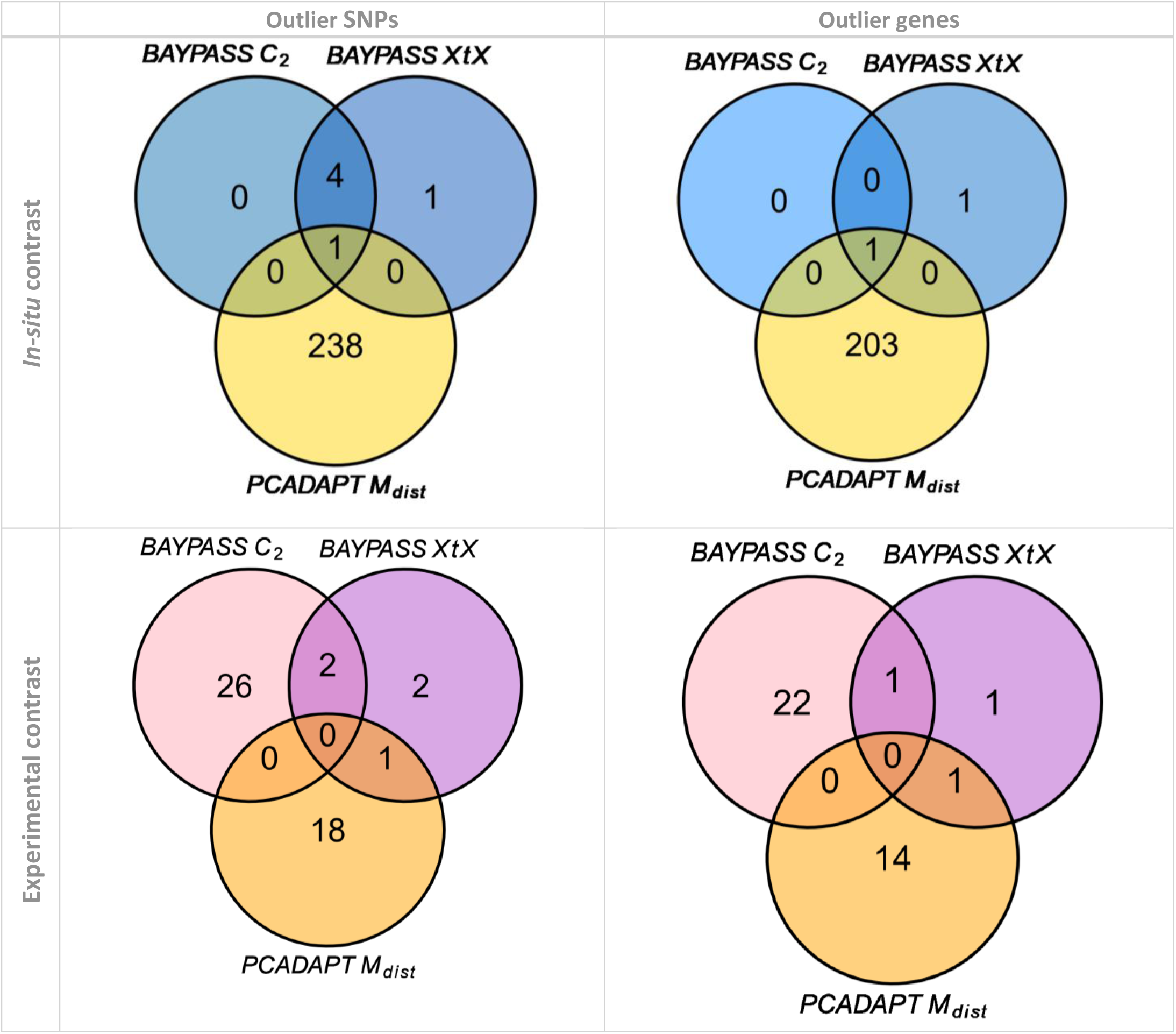
Venn diagrams illustrating the overlap between outlier SNPs and genes identified by *BAYPASS* and *PCAdapt*. We compare here three methods: *BAYPASS* based on *C2* statistics, *BAYPASS* based on *XtX* statistics and *PCAdapt* based on Mahalanobis distance (*Mdist*), for the two studied contrasts (top: *in-situ*; bottom: experimental) Plots on the left show the counts of outlier SNPs identified, while plots on the right show the counts of genes and intergenic regions containing these outlier SNPs.

**Figure S4.**
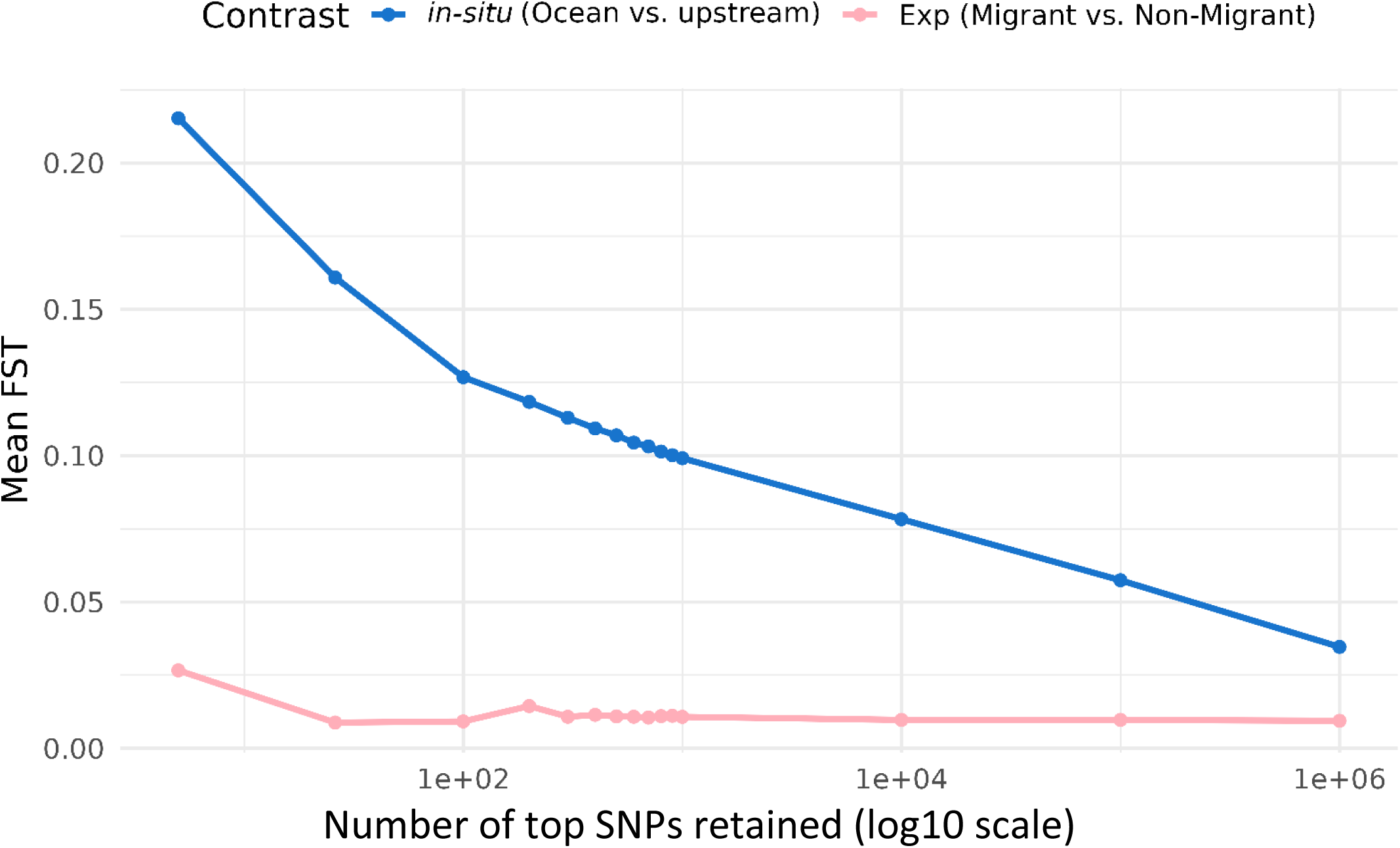
- Variation of mean *F_ST_* values computed for different sets of top *in-situ* SNPs. This figure is similar to Figure 3 in the main text, but includes a larger number of top- *in-situ*-SNP sets, displayed on a log-scaled x-axis. Blue lines correspond to average *F_ST_* between Ocean and Upstream sites, and pink lines to *F_ST_* between experimental Migrant and Non-migrant pools. We considered successive sets of SNPs identified from the *in-situ* contrast: the 5 “best” outliers, and then the top 26, 100, 200, …, 10^6^ SNPs with the lowest q-value.

**Figure S5.**
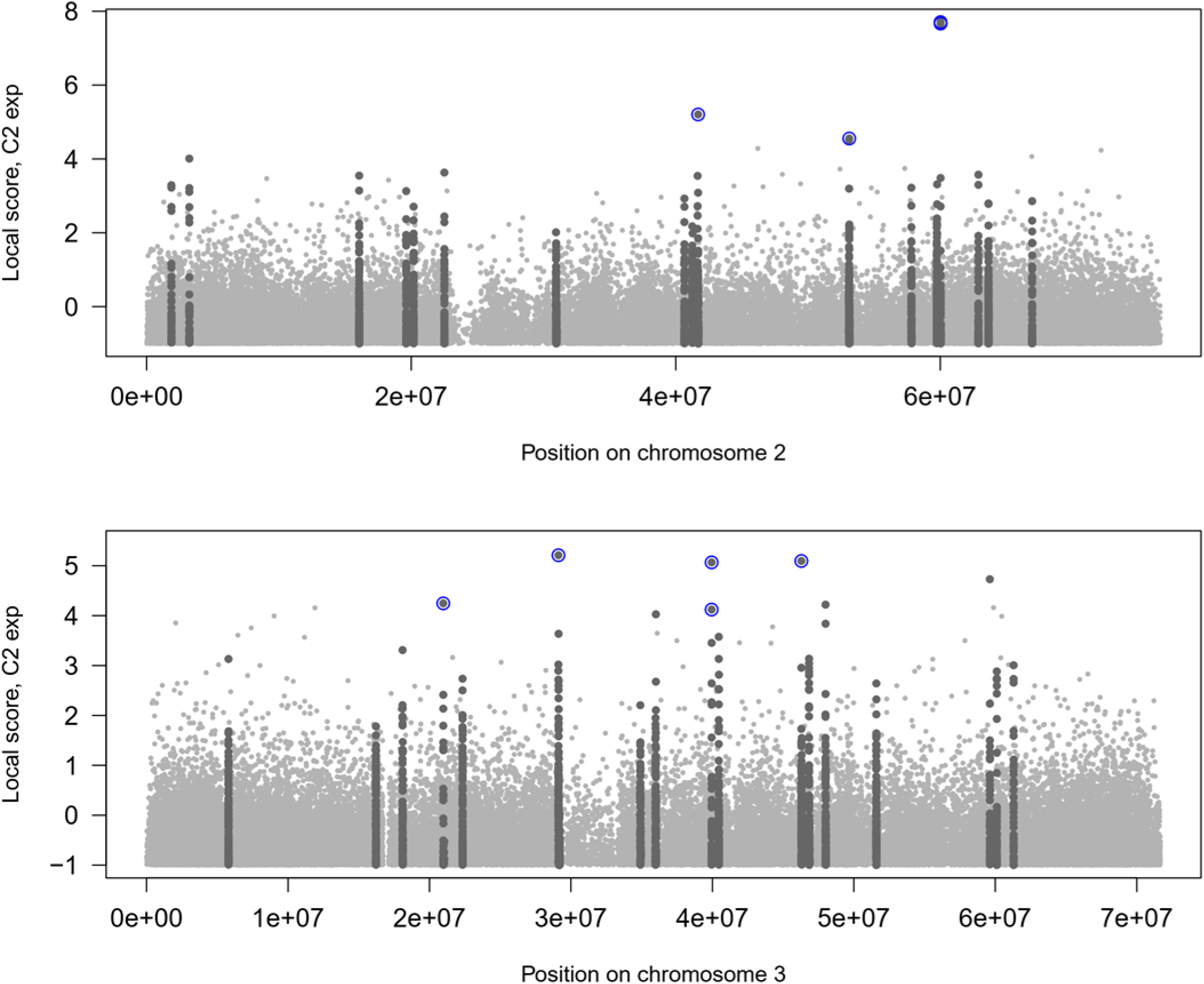
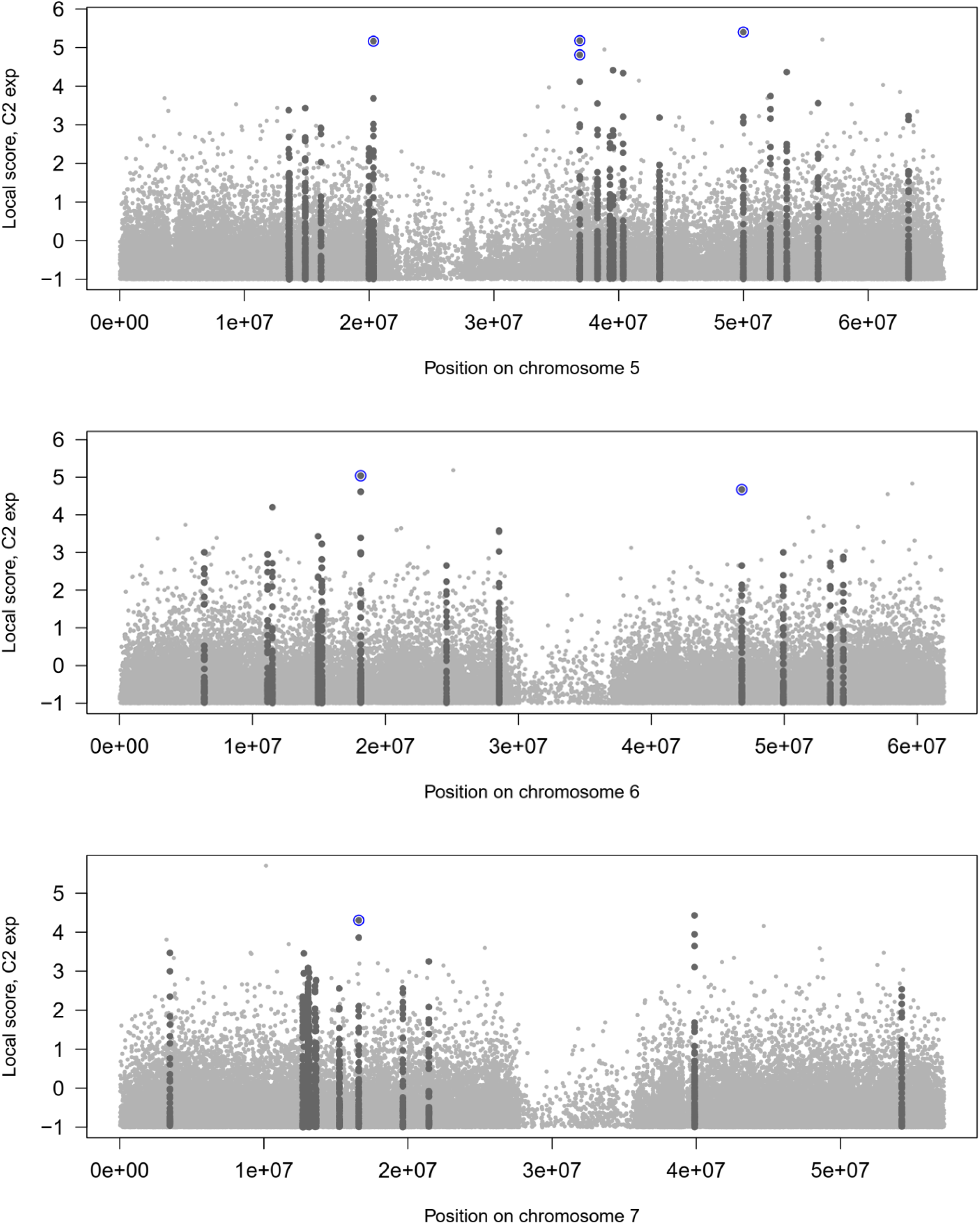
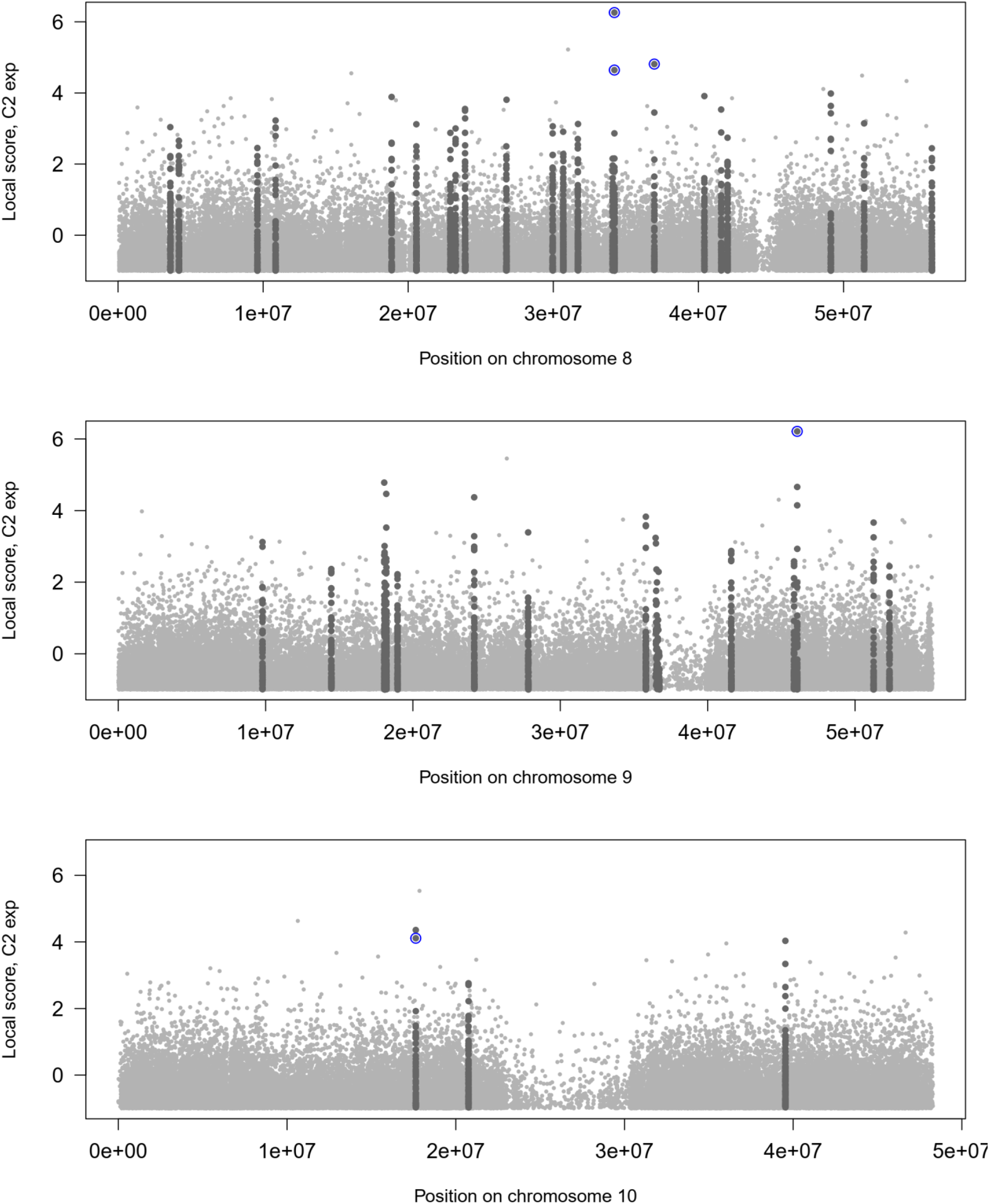
- Manhattan plots showing signatures of intragenerational selection detected with *BAYPASS* for the experimental contrast. Results are shown for 12 (chromosome 1 is shown in Figure 2, main text) of the 13 chromosomes containing best *F_ST_* outliers for differentiation between experimental Migrant and Non-migrant pools. The background local scores for all SNPs and the *C2_exp_* statistics appear in light grey points. Significant windows of high differentiation (Lindley score > 15) are marked by dark grey points. Blue circles highlight the best outliers SNPs, which passed the three filters (q-value, AFD, local score) described in Figure 1.

**Figure S6.**
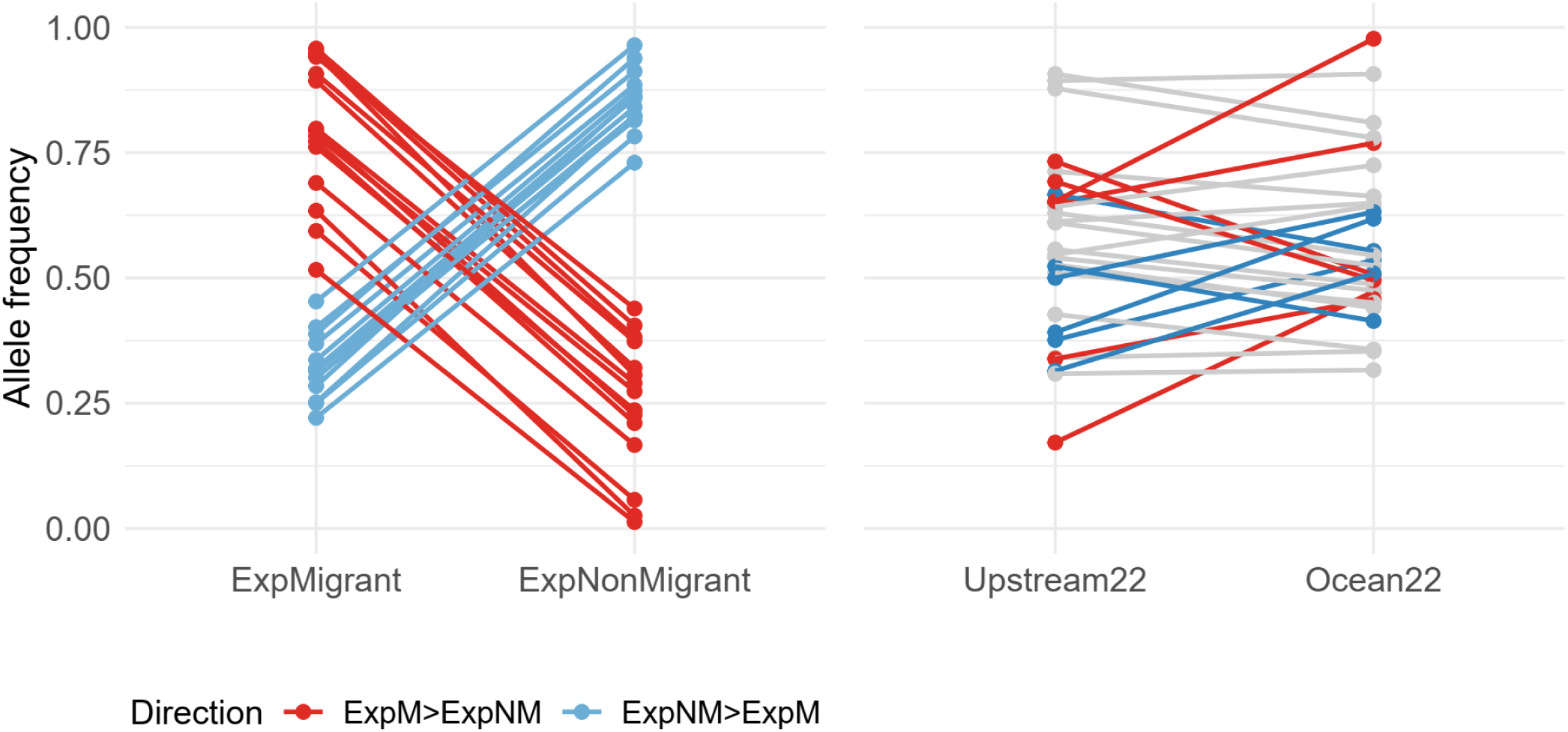
- Variation of allele frequencies at the 28 best experimental outlier SNPs. On the left, allele frequencies in each experimental pool (ExpMigrant, abbreviated ExpM vs ExpNonMigrant, abbreviated ExpNM) are shown for the 28 best outlier SNPs. Colours indicate the direction of allele frequency difference (AFD) between ExpM and ExpNM. For simplicity, this representation assumes that the proportion of reads carrying a given allele provides an unbiased estimate of allele frequency. The graph on the right shows allele frequencies for the same 28 outlier SNPs in the two in situ pools sampled in 2022, i.e. during the same months and year as the glass eels used in the experimental approach. Colours correspond to the direction of AFD observed in the experimental contrast. This graph shows that (1) 18 of the 28 top experimental outlier SNPs display negligible AFD (i.e. AFD < 0.10) between the Ocean and Upstream in situ locations (grey lines), and (2) among the 10 experimental outlier SNPs with non-negligible AFD (i.e. AFD > 0.10), the direction of allele-frequency change between Ocean and Upstream does not match the direction observed between ExpMigrant and ExpNonMigrant individuals

**Figure S7.**
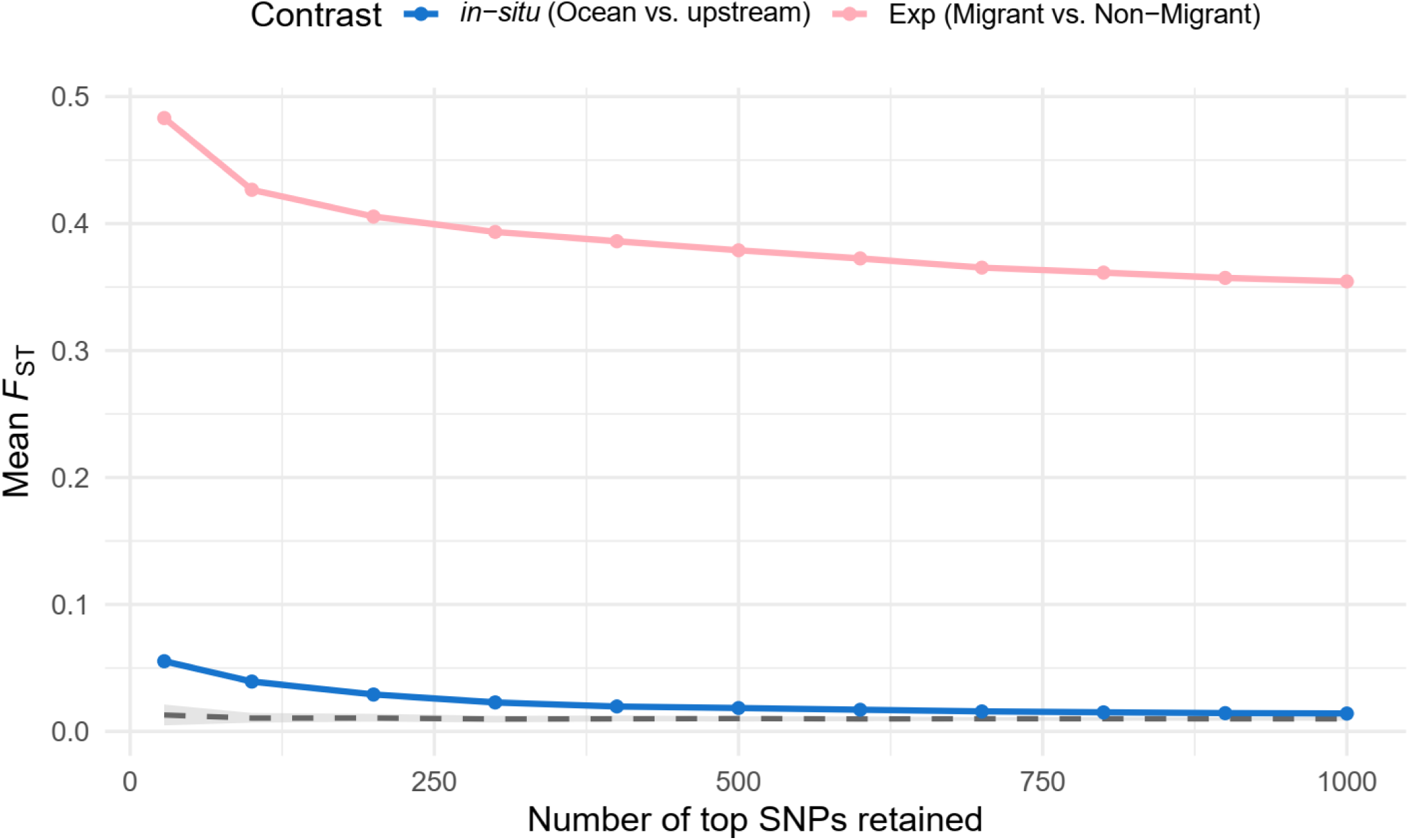
- Variation of mean *F_ST_* values computed for different sets of top experimental SNPs. This figure is similar to Figure 3 in the main text, but is based on top-SNP sets identified based on the experimental contrast (migrant vs non-migrant pools). The pink line corresponds to *F_ST_* between experimental Migrant and Non-migrant pools and the blue line to average *F_ST_* between Ocean and Upstream sites. The grey envelope represents the neutral expectation for *F_ST_* between experimental Migrant and Non-migrant pools; the dashed line indicates its median value We considered successive sets of SNPs identified from the experimental contrast: the 28 “best” outliers, and then the top 100, 200, …, 1000 SNPs with the lowest q-value. To build the neutral envelope, we randomly sampled, for each SNP set size, an equivalent number of loci with similar allele frequencies, and computed *F_ST_*. This was done 100 times to derive the 2.5–97.5% quantile range (grey area).

**Figure S8:**
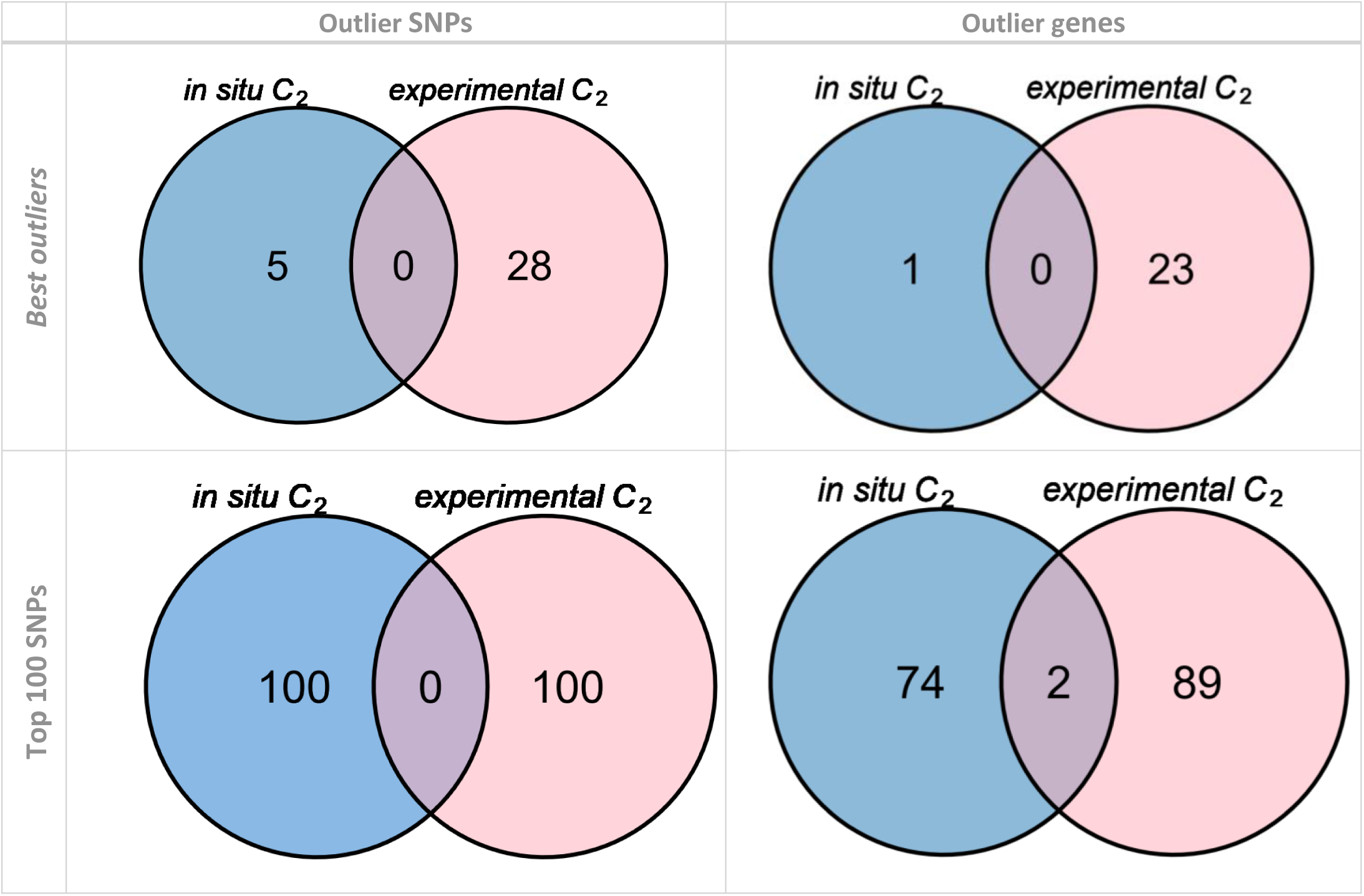
Venn diagrams of outlier SNPs and genes identified by the *in-situ* and experimental approaches. We compare here outlier SNPs and genes identified with *BAYPASS* based on *C2* statistics, for the two studied contrasts (blue: *in-situ*; pink: experimental). Plots on the left show the counts of outlier SNPs identified, while plots on the right show the counts of genes containing these outlier SNPs. On the bottom, diagram correspond to the best outlier SNPs, while on the bottom, we kept the top 100 SNPs identified for each contrast.

**Table S4:**
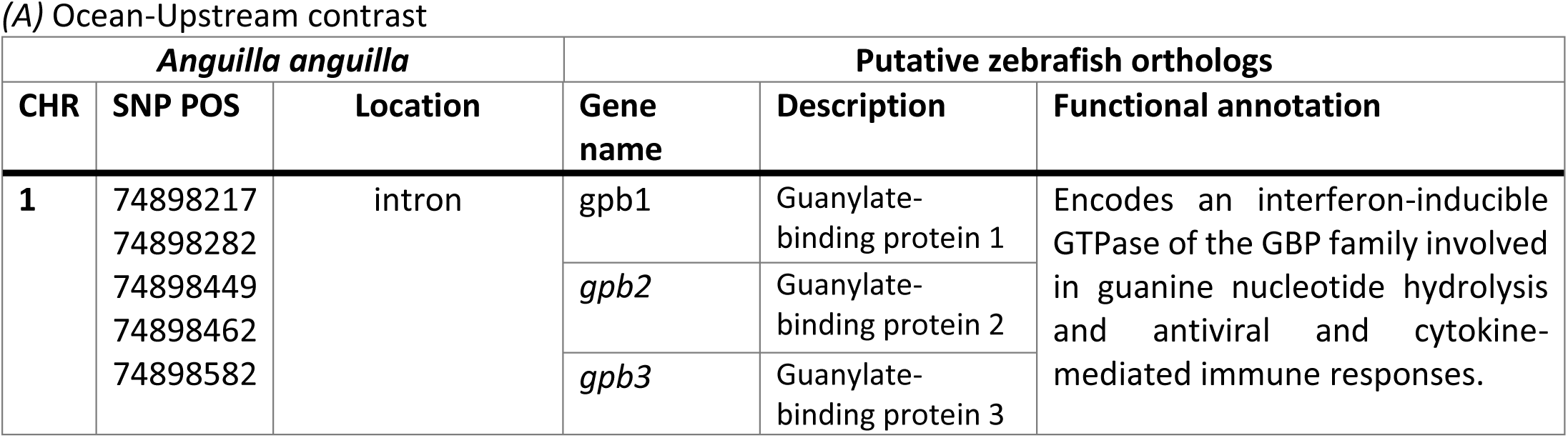
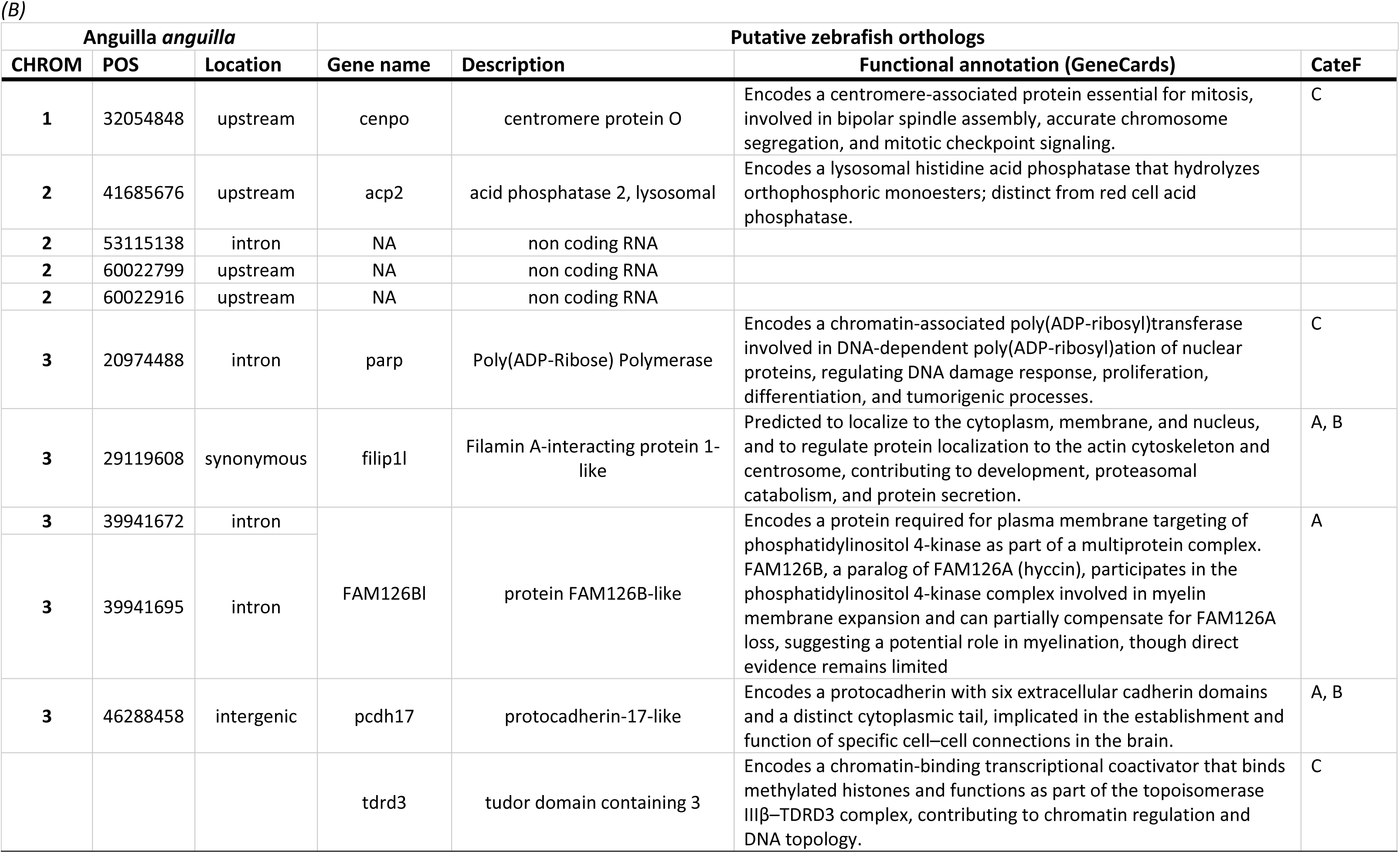
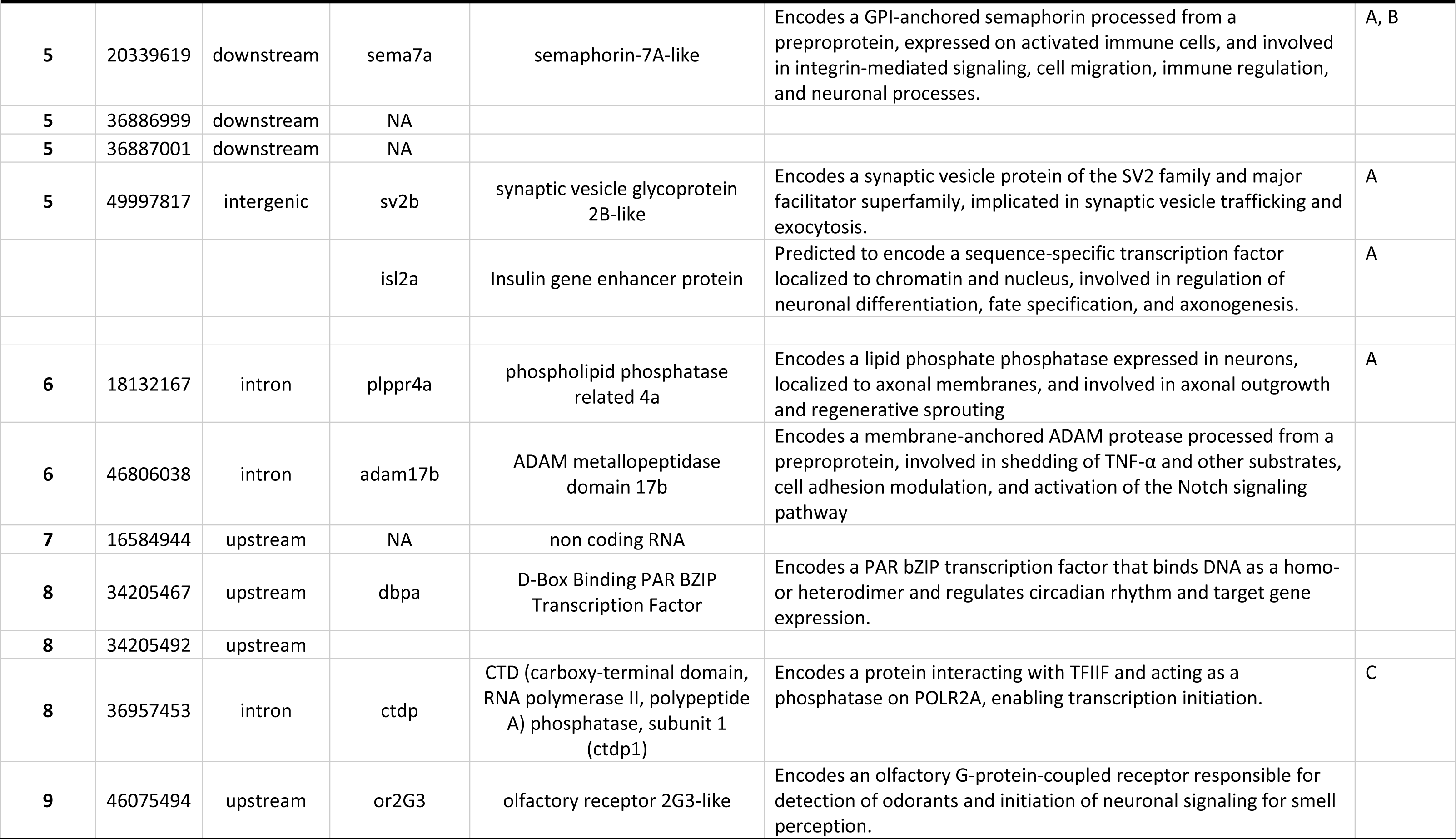
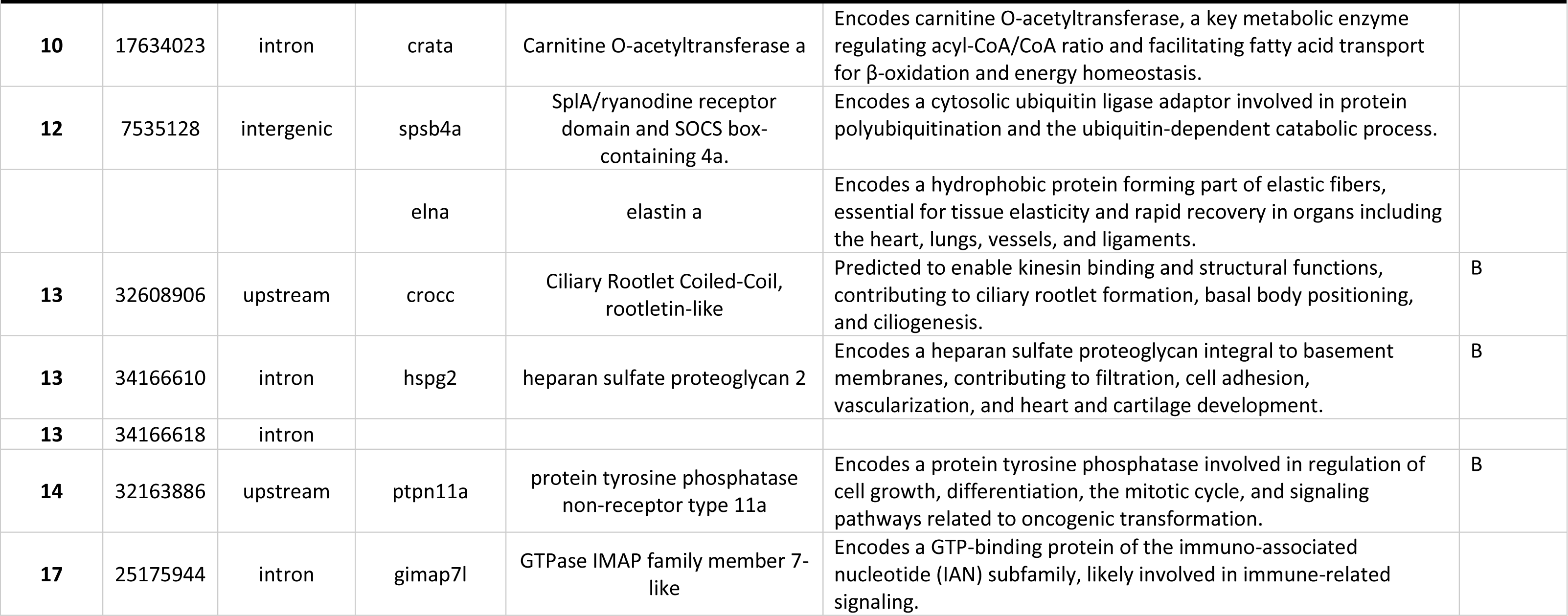
Functional annotation of the genes containing or close to outlier SNPs that differentiate (A) Ocean vs Upstream sites (in-situ contrast) and (B) Migrant vs non-migrant phenotypes (experimental contrast). For each SNP, we give the chromosome, the position, the location and eventual impact (synonymous within exon, intron, downstream or upstream the gen, intergenic), and information on the putative orthologs in zebrafish (gene name and description, unless the gene in uncharacterized- noted NA) as well as the functional annotation synthetized from GeneCards. For Table B, the last column (cateF) indicates the main categories of functional annotation to hich several genes here found to belong: A corresponds to « Neuronal development »; B corresponds to « Cell migration, cytoskeleton organization, tissue remodeling »; C corresponds to « Transcription / RNA processing ».

## Notes

### Competing Interest Statement

The authors have declared no competing interest.

### Summary of Updates

Minor corrections in the names of authors, previously mispelled, and the title of the manuscript.

